# JAK and MEK Pathways as Therapeutic Targets for Saphenous Vein Smooth Muscle Cell Dysfunction in Type 2 Diabetes *via* Regulation of Mitochondrial Activity

**DOI:** 10.1101/2025.10.17.682556

**Authors:** Israel O. Bolanle, Florah T. Moshapa, Gillian A. Durham, James P. Hobkirk, Kirsten Riches-Suman, Mahmoud Loubani, Roger G. Sturmey, Timothy M. Palmer

**Author notes:** Department of Bioengineering, Imperial College London, London SW7 2AZ, U.K. School of Pharmacy, Faculty of Health Sciences, University of Botswana, P/Bag-0022, Gaborone, Botswana (Florah T. Moshapa). Pears Cumbria School of Medicine, Carlisle CA1 2HH, U.K.

## Abstract

**Aims:** Glucose-driven mitochondrial dysfunction has been proposed to promote vascular cell proliferation and migration events responsible for saphenous vein graft failure (VGF) following bypass surgery. However, it is unclear how type 2 diabetes mellitus (T2DM) impacts mitochondrial function in human saphenous vein smooth muscle cells (HSVSMCs) responsible for the maladaptive remodelling responsible for VGF. Therefore, identifying and targeting the signalling pathways involved could offer new therapeutic options to limit VGF. Our aim was to identify signalling pathways that mediate any mitochondrial dysfunction in HSVSMCs *in vitro* and assess the impact of T2DM.

**Methods and results:** HSVSMCs explanted from surplus HSV tissues from consenting T2DM and non-diabetic patients undergoing coronary artery bypass graft surgery were treated with known activators and inhibitors of the JAK/STAT and MAPK/ERK pathways. Following this, real-time oxygen consumption rate (OCR) and extracellular acidification rate (ECAR) measures of mitochondrial function were then determined. Our findings revealed that both IL-6/sIL-6Rα trans-signalling complexes and PDGF-BB significantly increased OCR in HSVSMCs from T2DM patients but not non-diabetic controls. Meanwhile, only PDGF-BB increased ECAR in HSVSMCs from T2DM patients but not non-diabetic controls. The observed increases in OCR and ECAR were abolished by JAK1/2-selective inhibitor ruxolitinib. Furthermore, thrombin caused a significant increase in OCR specifically in HSVSMCs from T2DM patients, and this was abolished by MEK1/2-selective inhibitor trametinib. Both ruxolitinib and trametinib significantly reduced basal OCR and ECAR in HSVSMCs from both T2DM and non-diabetic patients.

**Conclusion:** Together, these findings demonstrate a JAK/STAT- and MAPK/ERK-mediated regulation of mitochondrial function in HSVSMCs. As such, they represent potential targets for regulation of HSVSMC function that can be explored for drug development to limit saphenous VGF in T2DM.

## 1. Introduction

Glucose-dependent alterations in cellular function have been suggested to promote proliferation and migration of vascular smooth muscle cells (VSMCs), which are linked to vein graft failure (VGF). ^1–3^ VSMCs from T2DM patients have distinct functional properties, ^5, 6^ including significantly greater migratory activity than those from non-diabetic controls. ^6, 7^ Additionally, type 2 diabetes mellitus (T2DM) is known to trigger changes in mitochondrial function ^4^ in VSMCs, and together, these findings hint at a link between T2DM, mitochondrial dysfunction, and VGF. Metabolic reprogramming, adaptive oxygen consumption and glucose utilisation are now appreciated as key mechanisms by which cells such as arterial endothelial and malignant cancer cells sustain high proliferative rates even in hostile environments in which the supplies of oxygen and essential nutrients are limited. ^8–10^ However, the impact of T2DM and possible downstream signalling pathways that regulate these processes in human saphenous vein smooth muscle cells (HSVSMCs), a crucial cell type implicated in the vascular dysfunction responsible for saphenous VGF, remain unclear. ^2, 3^

Mitochondrial oxygen consumption rate (OCR) and extracellular acidification rate (ECAR) have been described as gold standard measures for the determination of mitochondrial function in vascular endothelial cells (ECs), VSMCs, and platelets. ^11–13^ Since oxidative phosphorylation measured by OCR accounts for most of the increase in cellular ATP demand, the activity of this pathway is a major indicator of metabolic function. ^14, 15^ Measurement of OCR in the presence of certain inhibitors can offer valuable information on mitochondrial bioenergetic profiles. The components of oxygen consumption that are linked to ATP production can be determined by inhibitors that target the various complexes in the electron transport chain. ^15^ These components include the so-called coupled OCR, as well as the contribution made to OCR by passive or active proton leakage across the inner mitochondrial membrane, the difference between maximal oxygen consumption and basal (spare capacity), and non-mitochondrial oxygen consumption. Together, these tests offer a detailed picture of mitochondrial function. ^16^

In contrast, ECAR provides a measure of the lactic acid produced by cells whilst in culture. This is typically excreted by cells when lactate dehydrogenase converts pyruvate to lactate to replenish NAD+, which is necessary to sustain glycolysis. ^17, 18^ The flow through catabolic pathways necessary to produce ATP is, therefore, connected to a cell’s ECAR. The ECAR is also linked to ATP turnover, as the rates of ATP synthesis and consumption are equivalent in steady state. ^19^ Even though it is technically difficult to quantify ATP turnover, however, it may be demonstrated that fluctuations in extracellular fluxes are correlated with changes in ATP turnover rates. ^19^ For these reasons, mitochondrial OCR and ECAR are now appreciated as key parameters in assessing altered metabolic homeostasis in pathological conditions. ^11^

JAK/STAT and ERK 1/2 signalling pathways are important in numerous human physiological processes. ^20–22^ Importantly, activating these pathways with IL-6, PDGF, Ang II or thrombin can promote either vascular cell migration or proliferation events which are responsible for vascular remodelling. ^21–25^ However, their involvement in the modulation of mitochondrial function of vascular cells is poorly understood. Of importance is the fact that in patients with coronary artery disease (CAD) with or without T2DM, coronary artery bypass graft (CABG) using autologous saphenous vein remains the gold standard procedure for restoration of blood supply to the heart. ^26, 27^ Patients with T2DM who have CAD, however, are more susceptible to neointimal hyperplasia (NIH), vascular remodelling, and stenosis that result in VGF. ^28^ While the underlying mechanisms responsible for this are unclear, activities of the activators of the JAK/STAT and MAPK/ERK signalling pathways are upregulated in T2DM patients. ^23^ To prevent VGF, it may, therefore, be possible to find novel therapeutic targets by identifying characteristic metabolic modifications mediated by these downstream signalling pathways. Therefore, in this study, we assessed the role of JAK/STAT and MAPK/ERK in the regulation of mitochondrial functions (OCR and ECAR), along with the potential impact of T2DM on these processes in HSVSMCs *in vitro*.

## 2. Materials and Methods

**Table 2.1.**
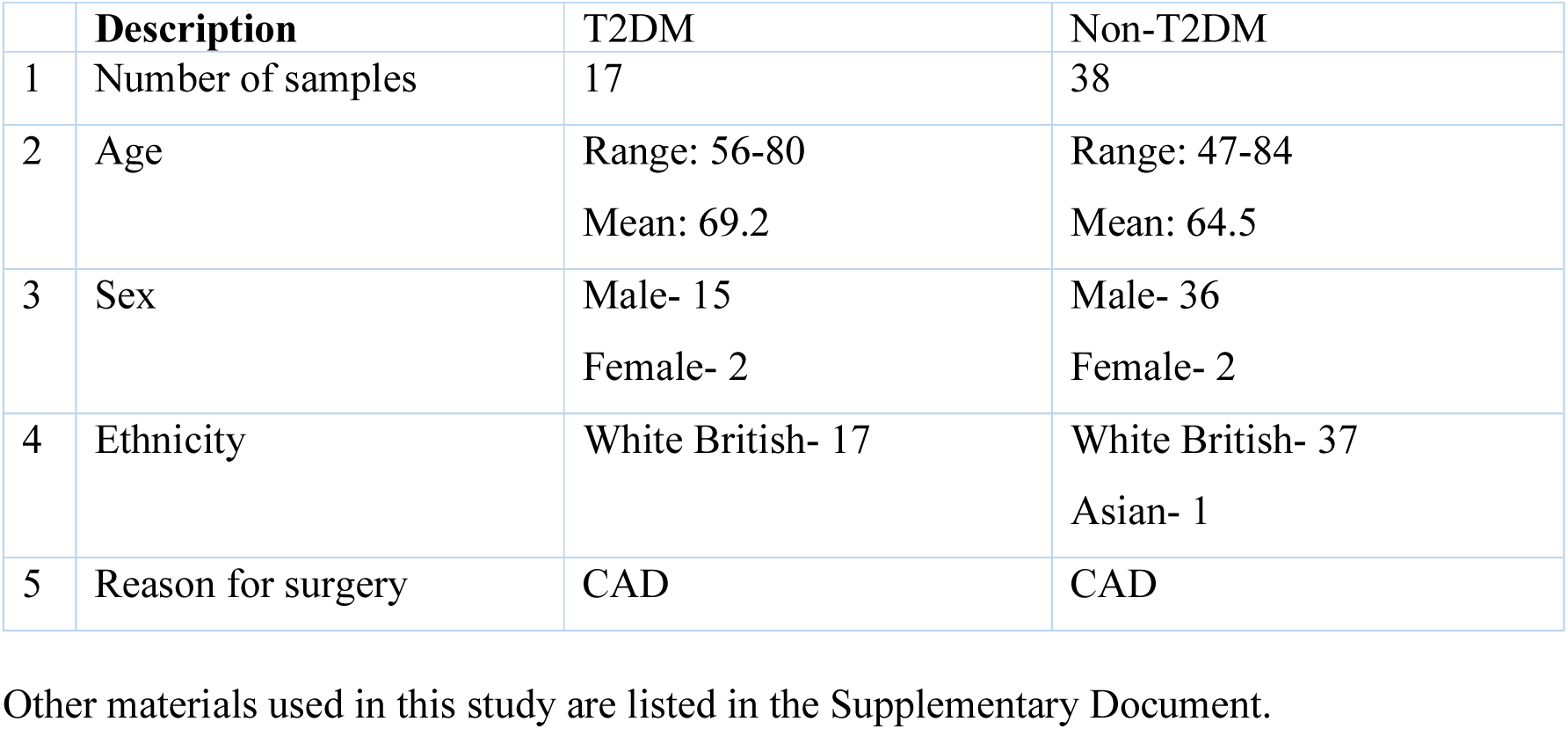
Patient information for HSV Samples. CAD: coronary artery disease; T2DM: type 2 diabetes mellitus. HSV samples were collected between January 2020 and May 2022 from patients undergoing elective coronary artery bypass graft procedure.

### 2.2 Methods

#### 2.2.1 Isolation of HSVSMCs

As shown in Table 1, surplus HSV tissues were obtained from consenting (written) T2DM and non-diabetic patients undergoing coronary artery bypass graft (CABG) surgery at the Hull University Teaching Hospitals NHS Trust Department of Cardiothoracic Surgery, Castle Hill Hospital, Cottingham, UK under UK Health Research Authority ethical approval (NHS REC:15/NE/0138). The procedures we used to obtain and process surplus HSV tissues adhered to the guidelines outlined in the 2024 revision of the Declaration of Helsinki.^29^ HSV tissues (length 0.5-4 cm) were stored at 4°C for a maximum of 48 hours in Dulbecco’s modified Eagle medium (DMEM) supplemented with 100 IU/ml penicillin, 100 µg/ml streptomycin, and 0.25 µg/ml fungizone. VSMCs were explanted from HSV as previously described. ^30^ Briefly, HSV tissue was placed in a 10 cm dish with enough medium to cover it, and a sterile blade was used to remove the perivascular fat, vein branches, connective tissue, and adventitia. The vein was then dissected longitudinally, and the endothelial and tunica intima layers gently removed with sterile surgical scissors and forceps. Then, HSV tissue was dissected into small fragments (∼1 mm^3^) which were transferred into a 25 cm^2^ tissue culture flask with 2 ml of SmGM2 and cultured at 37°C in 5% (v/v) CO_2_ in humidified atmosphere. Once HSVSMCs had migrated from the explant tissue, after approximately two weeks, the growth medium was topped up with 0.5 ml fresh SmGM2 twice weekly. Addition of 0.5 ml fresh SmGM2 continued until a total of 5 ml of growth medium had been added, then, 2.5 ml of growth medium was removed and replenished with fresh 2.5 ml SmGM2 until 90% confluence. Thereafter, HSVSMCs were transferred to a 75 cm^2^ tissue culture flask and fed with 10 ml of SmGM2 which was replaced twice weekly until cells reached 90% confluency and expanded for either cryopreservation in liquid nitrogen or downstream experimental applications. The integrity of HSVSMC reparations was assessed by immunofluorescence microscopy as previously described. ^31^

#### 2.2.2 HSVSMC mitochondrial stress analysis to determine OCR and ECARs

HSVSMCs were seeded in Seahorse XFp 8 well plates at a density of 10,000 cells per well and allowed to grow over 24 hrs. Following this, cells were treated with the respective inhibitors of the downstream signalling pathways. To evaluate the contribution of JAK/STAT signalling pathways, cells were treated with 0.1 μM JAK1/2-selective inhibitor ruxolitinib ^32^ for 90 min after which IL-6 (5 ng/ml) and sIL-6Rα (25 ng/ml) (IL-6/sIL-6Rα) or 10 ng/ml PDGF-BB was added for a further 24 hrs. To examine a role for ERK1,2 signalling pathways, cells were treated with 10 nM MEK1,2-selective inhibitor trametinib ^33^ for 90 min after which 100 nM Ang II or 1 U/ml thrombin were used to treat cells over 24 hrs in a cell culture incubator. Also, the probes of the XFp 8-well plate were hydrated overnight by adding 200 μl/well of Seahorse calibrant and 400 μl into the ridges of the plate to prevent evaporation. ^34^ A humidified incubator was used to incubate this hydrated sensor containing XFp 8 well plates overnight at 37°C.

On the day of assay, modulators of mitochondrial respiration; 20 μl oligomycin (1.5 μM), 22 μl carbonyl cyanide p-(trifluoromethoxy) phenylhydrazone (FCCP) (5 μM), and 25 μl rotenone (5 μM) and antimycin A (5 μM) complex were added to the injection ports of the pre-hydrated Seahorse XFp 8 well plate. ^34^ The Seahorse XFp 8 well plate was kept in the non-CO_2_ incubator for 30 minutes to allow the drugs to equilibrate. Each treatment was done in duplicate. Following this, plates were mounted in the Agilent Seahorse XFp Analyzer (model number 102745-100) and allowed to calibrate. While calibrating, the growth media on the pre-treated cells were withdrawn and 180 μl of the XF DMEM medium was added to each well. Following calibration, the cells were mounted in the Agilent Seahorse XFp Analyzer (model number 102745-100) and real-time OCR and ECAR of the pre-treated cells were determined. ^34^ After assay, treated cells were lysed and the protein concentrations were determined using a BCA assay. Four biological replicates using samples from different patients were performed. Values for OCR and ECAR were normalised to protein concentration.

#### 2.2.4 Real-time quantitative PCR

qPCR was used to determine mtDNA copy number of mitochondrially-encoded gene for cytochrome c oxidase subunit I (COI), a key protein in the mitochondrial respiratory chain. ^35, 36^ A total volume of 20 μl PCR mixture, comprising of 1 ng of cDNA, 10 μl of SYBR Green Master Mix, 1 μl of the appropriate primer, and 7.4 μl of RNase-free H_2_O was added to each well of a 96-well PCR plate. The 96-well plate was then installed in the StepOne machine after being sealed with optical adhesive film. StepOne software was used to perform the real-time quantitative PCR (qPCR) following the manufacturer’s recommendations. The cycle conditions for real-time qPCR were optimised and set at 95°C for 10 minutes, followed by 35 cycles of 95°C for 30 seconds, 57°C for 30 seconds, and 72°C for 30 seconds. Every assay was carried out twice, and the average was determined. RNase-free water served as the negative control. Each sample’s average C_T_ values were computed and normalised to 18S RNA controls.

**Table 2.2.**
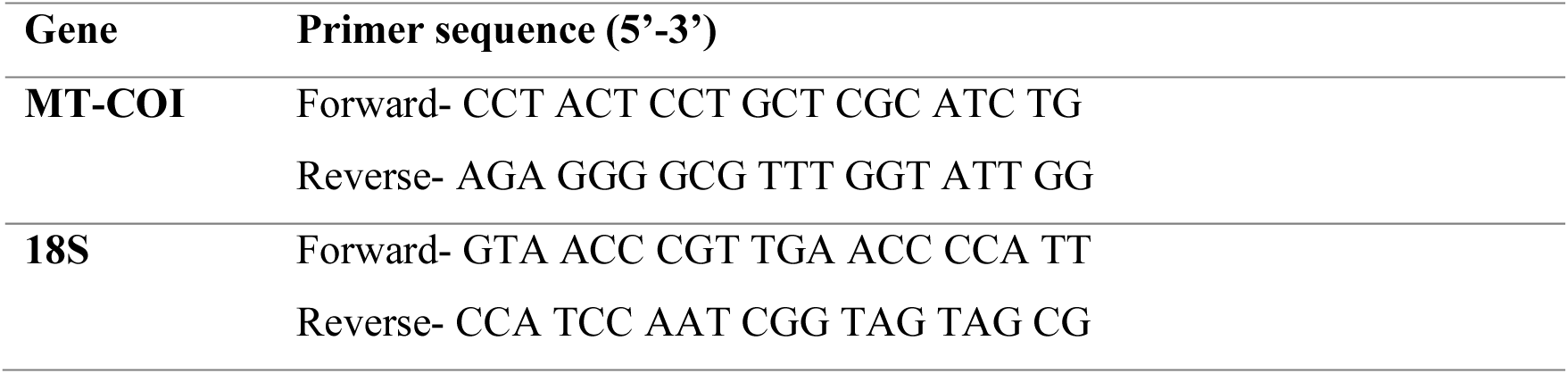
PCR primer sequence.

#### 2.2.5 Statistical analysis

Statistical analysis was carried out using GraphPad Prism® 6, San Diego, USA. Data are presented as mean ± standard error of mean (S.E.M) and were analysed using an independent t-test and one-way analysis of variance, followed by Dunnett’s post-hoc test to determine significant difference between means where *p*<0.05 was considered statistically significant.

## 3. Results

### 3.1. Effect of IL-6/sIL-6Rα on OCR in HSVSMCs from T2DM and non-diabetic patients

Studies have shown that T2DM is associated with elevated levels of pleiotropic cytokines such as IL-6, which can cause a number of adverse cellular effects, including mitochondrial dysfunction. ^23^ However, the mechanism(s) and impact on mitochondrial function of VSMCs are unknown. As shown in Figure 1A, an IL-6/sIL-6Rα trans-signalling complex was found to significantly increase OCR in HSVSMCs from T2DM patients at maximal respiration (*p*<0.05 versus unstimulated cells, n=4), but not in those from non-diabetic HSVSMCs (Figure 1B). The IL-6/sIL-6Rα-stimulated increase in OCR found in HSVSMCs from T2DM patients was abolished by pre-treatment with JAK1/2-selective inhibitor ruxolitinib (Figure 1A), at a concentration (0.1 μM) sufficient to significantly inhibit IL-6/sIL-6Rα-mediated phosphorylation of STAT3 on Tyr705 (Supplementary Information Figure S1). Furthermore, ruxolitinib significantly decreased OCR in HSVSMCs from non-diabetic patients at basal and maximal respiration with or without IL-6/sIL-6Rα stimulation (*p*<0.05 versus unstimulated cells, n=4; Figure 1B). No significant differences in OR between HSVSMCs from T2DM and non-diabetic patients were observed (Figure 1C).

**Figure 1:**
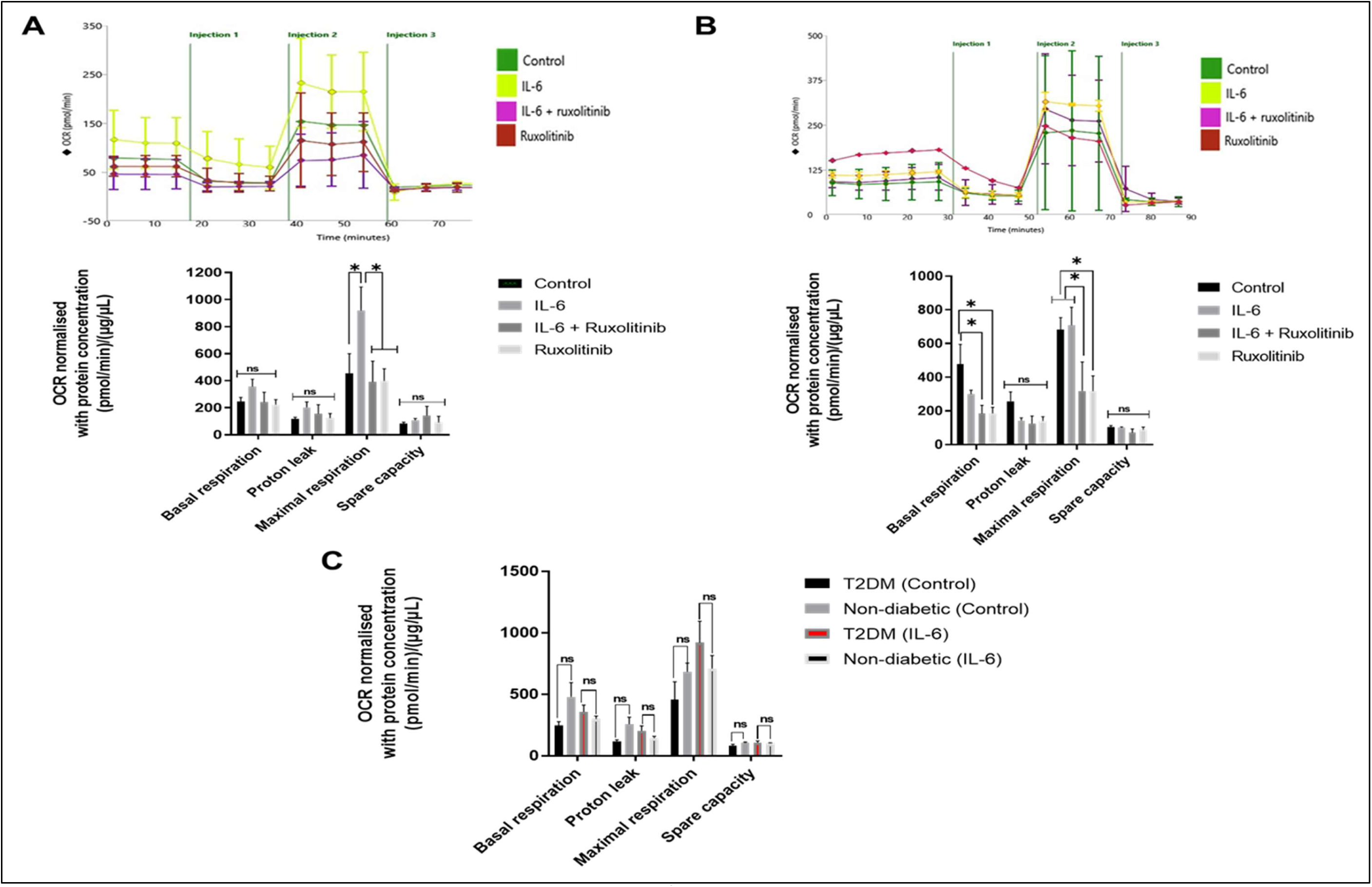
Mitochondria stress analysis to determine the OCR of HSVSMCs from T2DM and non-T2DM patients after stimulation with IL-6/sIL-6Rα +/- ruxolitinib. **(A) Upper panel:** representative time course curve of OCR of HSVSMCs from T2DM patients treated with IL-6/sIL-6Rα (IL-6 (5 ng/ml) and sIL-6Rα (25 ng/ml)) +/- ruxolitinib (0.1 μM) and untreated control. **Lower panel:** comparative analysis of normalised peak OCR between treatment groups and untreated control at basal glycolysis, and after addition of inhibitors of mitochondrial respiration and the uncoupler. Normalised data are presented as mean ± SEM from n=4 biological replicates using HSVSMC samples from different T2DM patients. **(B) Upper panel:** representative time course curve of OCR of HSVSMCs from non-diabetic patients treated with IL-6/sIL-6Rα (IL-6 (5 ng/ml) and sIL-6Rα (25 ng/ml)) +/- ruxolitinib (0.1 μM) and untreated control. **Lower panel:** comparative analysis of normalised peak OCR between treatment groups and untreated control at basal glycolysis, and after addition of inhibitors of mitochondrial respiration and the uncoupler. Normalised data are presented as mean ± SEM from n=4 biological replicates using HSVSMC samples from different non-diabetic patients. **(C)** Comparison of normalised peak OCR of unstimulated and IL-6/sIL-6Rα-stimulated HSVSMCs from T2DM patients versus non-diabetic control. Normalised data are presented as mean ± SEM and statistical significance was assessed using an independent t-test, n=4 biological replicates using HSVSMC samples from different T2DM and non-diabetic patients. In the lower panels of Figures A and B, normalised data from different treatment conditions were compared with data from the normalised untreated control. Statistical significance was assessed using one-way ANOVA followed by Dunnett’s post-hoc test to determine significant differences between means, **\****P* < 0.05. IL-6: IL-6/sIL-6Rα; OCR: oxygen consumption rate.

### 3.2. Effect of IL-6/sIL-6Rα on the ECAR in HSVSMCs from T2DM and non-diabetic patients

The effect of pro-inflammatory cytokines such as IL-6 on ECAR of vascular cells is currently unclear. ECAR allows for direct quantification of glycolysis. ^17^ We therefore determined the ECAR of HSVSMCs from T2DM and non-diabetic patients after treatment with IL-6/sIL-6Rα following pre-treatment with or without ruxolitinib. Our findings suggest that IL-6/sIL-6Rα did not cause any significant change in the ECAR of HSVSMCs from either T2DM or non-diabetic patients (Figures 2A and 2B). However, with or without IL-6/sIL-6Rα stimulation, ruxolitinib significantly (*p*<0.05 versus unstimulated cells, n=4) decreased ECAR at basal and maximal glycolysis, and (*p*<0.01 versus unstimulated cells, n=4) at glycolytic reserve in HSVSMCs of non-diabetic patients (Figure 2B). No significant differences in the ECAR of HSVSMCs from T2DM and non-diabetic patients after stimulation with IL-6/sIL-6R were observed (Figure 2C).

**Figure 2:**
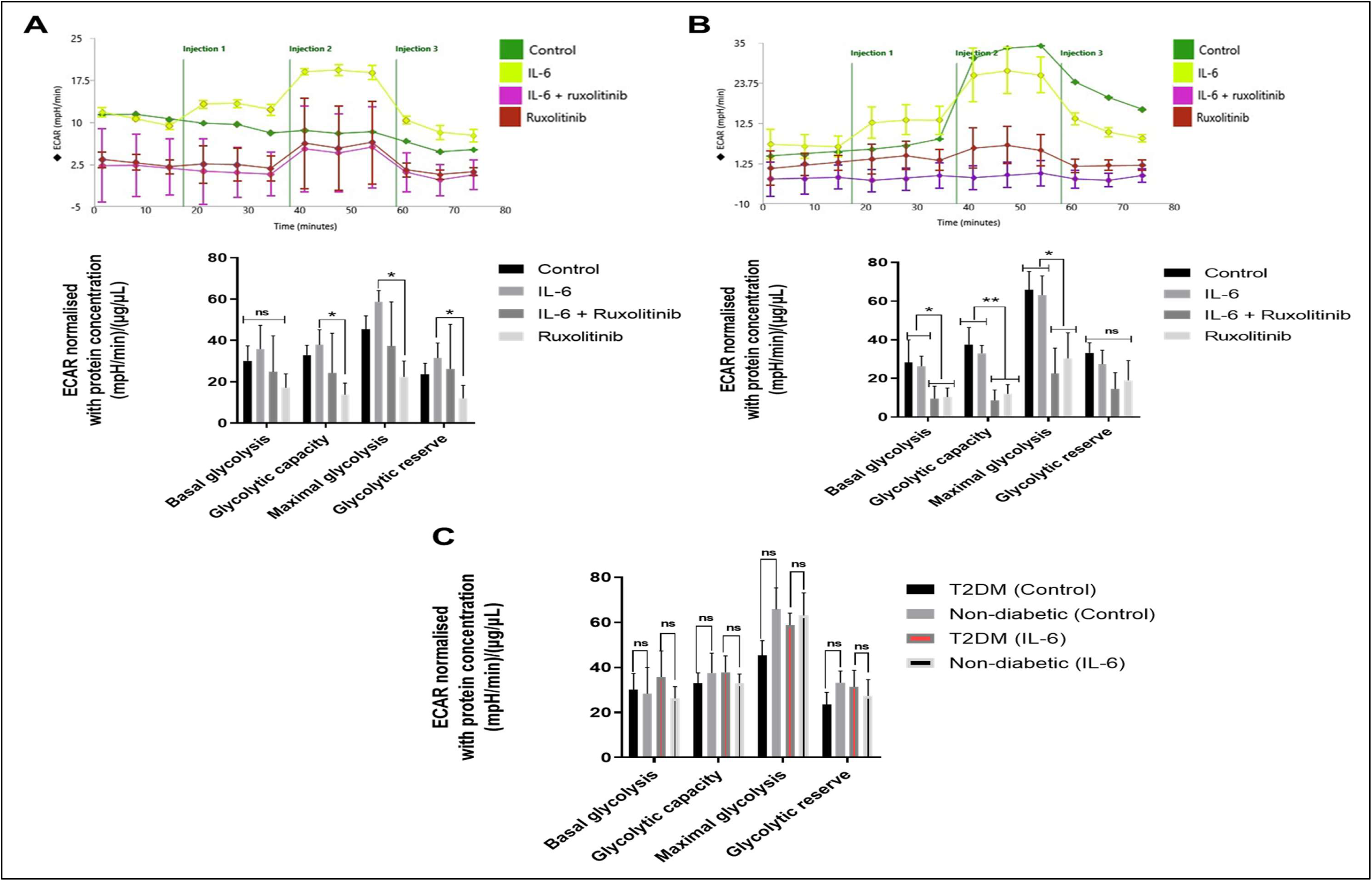
Mitochondria stress analysis to determine the ECAR of HSVSMCs from T2DM and non-T2DM patients after stimulation with IL-6/sIL-6Rα +/- ruxolitinib. **(D) Upper panel:** representative time course curve of ECAR of HSVSMCs from T2DM patients treated with IL-6/sIL-6Rα (IL-6 (5 ng/ml) and sIL-6Rα (25 ng/ml)) +/- ruxolitinib (0.1 μM) and untreated control. **Lower panel:** comparative analysis of normalised peak ECAR between treatment groups and untreated control at basal glycolysis, and after addition of inhibitors of mitochondrial respiration and the uncoupler. Normalised data are presented as mean ± SEM from n=4 biological replicates using HSVSMC samples from different T2DM patients. **(E) Upper panel:** representative time course curve of ECAR of HSVSMCs from non-diabetic patients treated with IL-6/sIL-6Rα (IL-6 (5 ng/ml) and sIL-6Rα (25 ng/ml)) +/- ruxolitinib (0.1 μM) and untreated control. **Lower panel:** comparative analysis of normalised peak ECAR between treatment groups and untreated control at basal glycolysis, and after addition of inhibitors of mitochondrial respiration and the uncoupler. Normalised data are presented as mean ± SEM from n=4 biological replicates using HSVSMC samples from different non-diabetic patients. **(F)** Comparison of normalised peak ECAR of unstimulated and IL-6/sIL-6Rα-stimulated HSVSMCs from T2DM patients versus non-diabetic control. Normalised data are presented as mean ± SEM and statistical significance was assessed using an independent t-test, n=4 biological replicates using HSVSMC samples from different T2DM and non-diabetic patients. In the lower panels of Figures A and B, normalised data from different treatment conditions were compared with data from the normalised untreated control. Statistical significance was assessed using one-way ANOVA followed by Dunnett’s post-hoc test to determine significant differences between means, **\****P* < 0.05; \*\**P* < 0.01. IL-6: IL-6/sIL-6Rα; ECAR: extracellular acidification rate.

### 3.3. Effect of PDGF-BB on the OCR in HSVSMCs from T2DM and non-diabetic patients

Given that the OCR is proportional to mitochondrial respiration, ^17^ it is unclear how growth factors like PDGF alter this cellular metabolic index (OCR) in HSVSMCs. To investigate this, the OCRs of HSVSMCs from T2DM patients and non-diabetic controls were assessed after treatment with PDGF-BB in the presence or absence of ruxolitinib. As shown in Figure 3A, PDGF-BB significantly increased OCR in HSVSMCs from T2DM patients (*p*<0.05 versus unstimulated cells, n=4) and maximum respiration (*p*<0.01 versus unstimulated cells, n=4). However, PDGF-BB had no significant effect on the OCR of HSVSMCs from non-diabetic patients (Figure 3B). This increase in OCR in T2DM patient HSVSMCs was abolished by pre-treatment with ruxolitinib (Figure 3A). Ruxolitinib also significantly reduced OCR at maximum respiration (*p*<0.05 versus unstimulated cells and *p*<0.01 versus PDGF-BB-stimulated cells, n=4) of HSVSMCs from T2DM patients (Figure 3A); a pattern also seen in HSVSMCs from non-diabetic controls (Figures 3B) (*p*<0.05 versus unstimulated and PDGF-BB stimulated, n=4). In contrast there were no significant differences between OCRs in HSVSMCs from T2DM and non-diabetic patients (Figure 3C).

**Figure 3:**
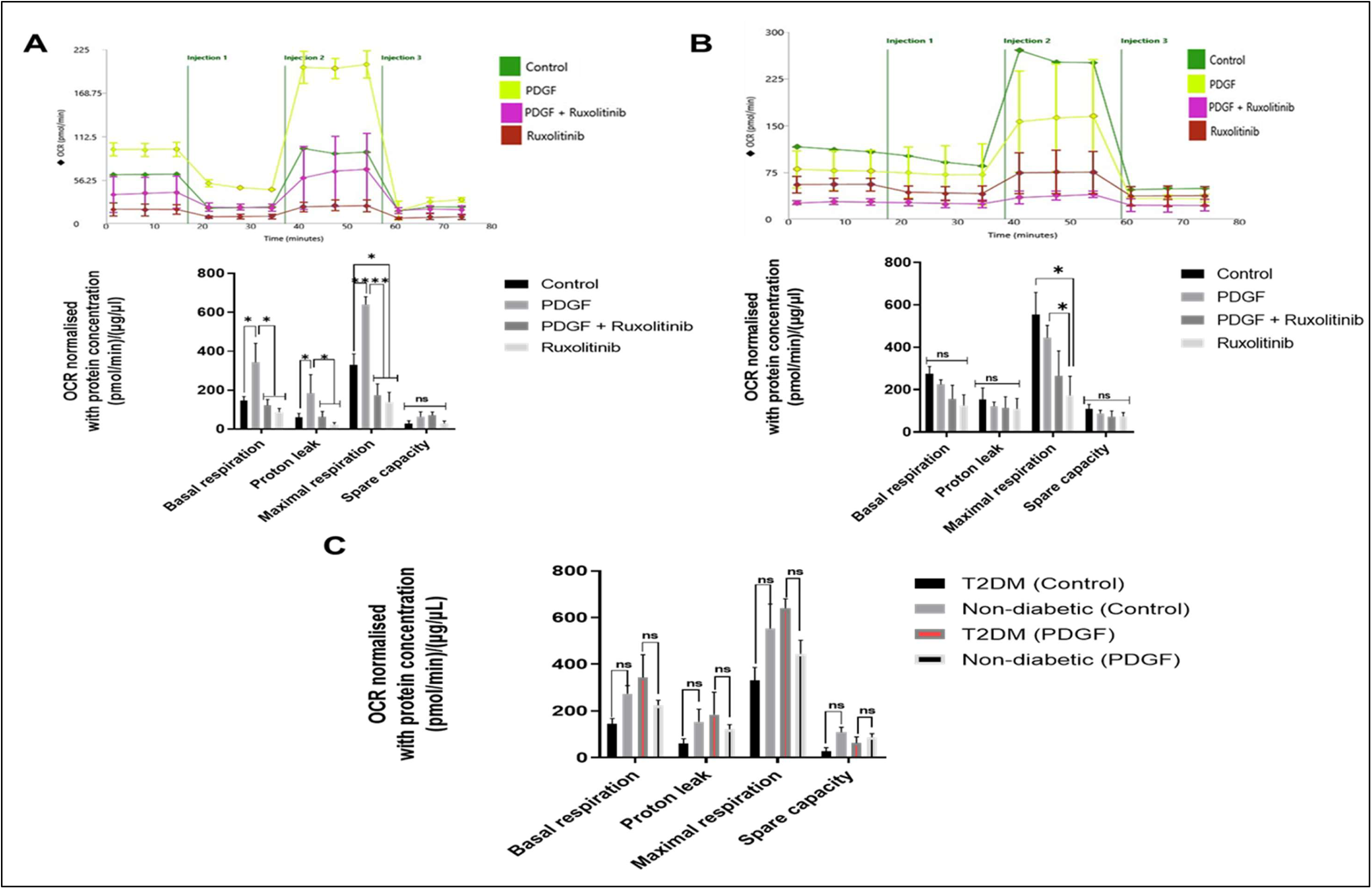
Mitochondria stress analysis to determine the OCR of HSVSMCs from T2DM and non-T2DM patients after stimulation with PDGF-BB +/- ruxolitinib. **(G) Upper panel:** representative time course curve of OCR of HSVSMCs from T2DM patients treated with PDGF-BB (10 ng/ml) +/- ruxolitinib (0.1 μM) and untreated control. **Lower panel:** comparative analysis of normalised peak OCR between treatment groups and untreated control at basal glycolysis, and after addition of inhibitors of mitochondrial respiration and the uncoupler. Normalised data are presented as mean ± SEM from n=4 biological replicates using HSVSMC samples from different T2DM patients. **(H) Upper panel:** representative time course curve of OCR of HSVSMCs from non-diabetic patients treated with PDGF-BB (10 ng/ml) +/- ruxolitinib (0.1 μM) and untreated control. **Lower panel:** comparative analysis of normalised peak OCR between treatment groups and untreated control at basal glycolysis, and after addition of inhibitors of mitochondrial respiration and the uncoupler. Normalised data are presented as mean ± SEM from n=4 biological replicates using HSVSMC samples from different non-diabetic patients. **(I)** Comparison of normalised peak OCR of unstimulated and PDGF-BB-stimulated HSVSMCs from T2DM patients versus non-diabetic control. Normalised data are presented as mean ± SEM and statistical significance was assessed using an independent t-test, n=4 biological replicates using HSVSMC samples from different T2DM and non-diabetic patients. In the lower panels of Figures A and B, normalised data from different treatment conditions were compared with data from the normalised untreated control. Statistical significance was assessed using one-way ANOVA followed by Dunnett’s post-hoc test to determine significant differences between means, **\****P* < 0.05; \*\**P* < 0.01. PDGF: PDGF-BB; OCR: oxygen consumption rate.

### 3.4. Effect of PDGF-BB on the ECAR in HSVSMCs from T2DM and non-diabetic patients

We next determined the ECAR of HSVSMCs from T2DM and non-diabetic control treated with or without PDGF-BB following pre-treatment with or without ruxolitinib. As shown in Figure 4A, PDGF-BB significantly increased basal ECAR, as well as glycolytic capacity, maximal glycolysis, and glycolytic reserve in HSVSMCs from T2DM patients (*p*<0.05 versus unstimulated cells, n=4) but not in non-diabetic controls (Figure 4B). Furthermore, ruxolitinib significantly decreased basal ECAR, glycolytic capacity, maximal glycolysis, and glycolytic reserve in HSVSMCs from non-diabetic patients (*p*<0.05 versus unstimulated and PDGF-BB-stimulated cells but not in those from T2DM patients (Figure 4B). Direct comparison of the ECAR of HSVSMCs from T2DM versus non-diabetic patients also revealed a significantly higher ECAR in HSVSMCs from T2DM compared with non-diabetic controls (*p*<0.05 n=4; Figure 4C) at maximal glycolysis.

**Figure 4:**
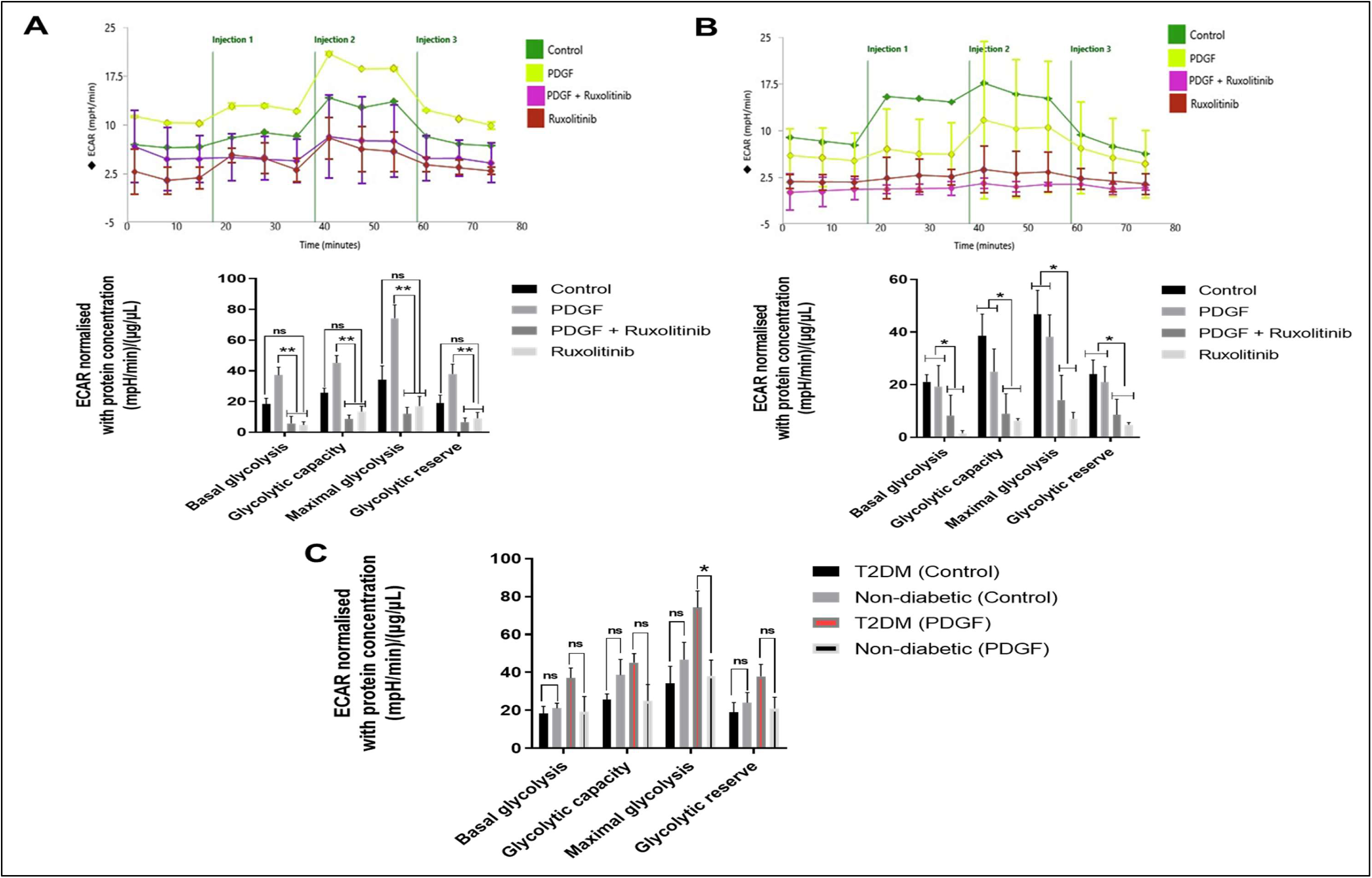
Mitochondria stress analysis to determine the ECAR of HSVSMCs from T2DM and non-T2DM patients after stimulation with PDGF-BB +/- ruxolitinib. **(J) Upper panel:** representative time course curve of ECAR of HSVSMCs from T2DM patients treated with PDGF-BB (10 ng/ml) +/- ruxolitinib (0.1 μM) and untreated control. **Lower panel:** comparative analysis of normalised peak ECAR between treatment groups and untreated control at basal glycolysis, and after addition of inhibitors of mitochondrial respiration and the uncoupler. Normalised data are presented as mean ± SEM from n=4 biological replicates using HSVSMC samples from different T2DM patients. **(K) Upper panel:** representative time course curve of ECAR of HSVSMCs from non-diabetic patients treated with PDGF-BB (10 ng/ml) +/- ruxolitinib (0.1 μM) and untreated control. **Lower panel:** comparative analysis of normalised peak ECAR between treatment groups and untreated control at basal glycolysis, and after addition of inhibitors of mitochondrial respiration and the uncoupler. Normalised data are presented as mean ± SEM from n=4 biological replicates using HSVSMC samples from different non-diabetic patients. **(L)** Comparison of normalised peak ECAR of unstimulated and PDGF-BB-stimulated HSVSMCs from T2DM patients versus non-diabetic control. Normalised data are presented as mean ± SEM and statistical significance was assessed using an independent t-test, n=4 biological replicates using HSVSMC samples from different T2DM and non-diabetic patients. In the lower panels of Figures A and B, normalised data from different treatment conditions were compared with data from the normalised untreated control. Statistical significance was assessed using one-way ANOVA followed by Dunnett’s post-hoc test to determine significant differences between means, **\****P* < 0.05; \*\**P* < 0.01. PDGF: PDGF-BB; ECAR: extracellular acidification rate.

### 3.4. mtDNA copy number in HSVSMCs from T2DM and non-diabetic patients after treatments with IL-6/sIL-6Rα and PDGF-BB +/-ruxolitinib

As JAK/STAT activators IL-6/sIL-6Rα and PDGF-BB caused a significant increase in the OCR of HSVSMCs from T2DM (Figures 1A and 3A) but not non-diabetic patients (Figures 1B and 3B), it was therefore necessary to ascertain whether this increase was associated with a parallel increase in mitochondria. Therefore, qPCR of mitochondrially-encoded gene for COI was used to determine mtDNA copy number.

As shown in Figure 5A, there was no significant difference in the mtDNA copy number in HSVSMCs from T2DM patients after treatments with IL-6/sIL-6Rα+/-ruxolitinib compared with untreated control. Similarly, in the HSVSMCs from non-diabetic patients, there was no significant difference in the mtDNA copy number after treatment with IL-6/sIL-6Rα+/- ruxolitinib compared with untreated control (Figure 5B). Comparison between HSVSMCs from T2DM patients versus non-diabetic control treated with IL-6/sIL-6Rα+/-ruxolitinib showed no significant difference (Figure 5C). Furthermore, as shown in Figure 5D, there was no significant difference in the mtDNA copy number in HSVSMCs from T2DM patients after treatments with PDGF-BB+/-ruxolitinib compared with untreated control. Similarly, in the HSVSMCs from non-diabetic patients, there was no significant difference in the mtDNA copy number after treatment with PDGF-BB+/-ruxolitinib compared with untreated control (Figure 5E). Also, comparison between HSVSMCs from T2DM patients versus non-diabetic control treated with PDGF-BB+/-ruxolitinib showed no significant difference (Figure 5F).

**Figure 5:**
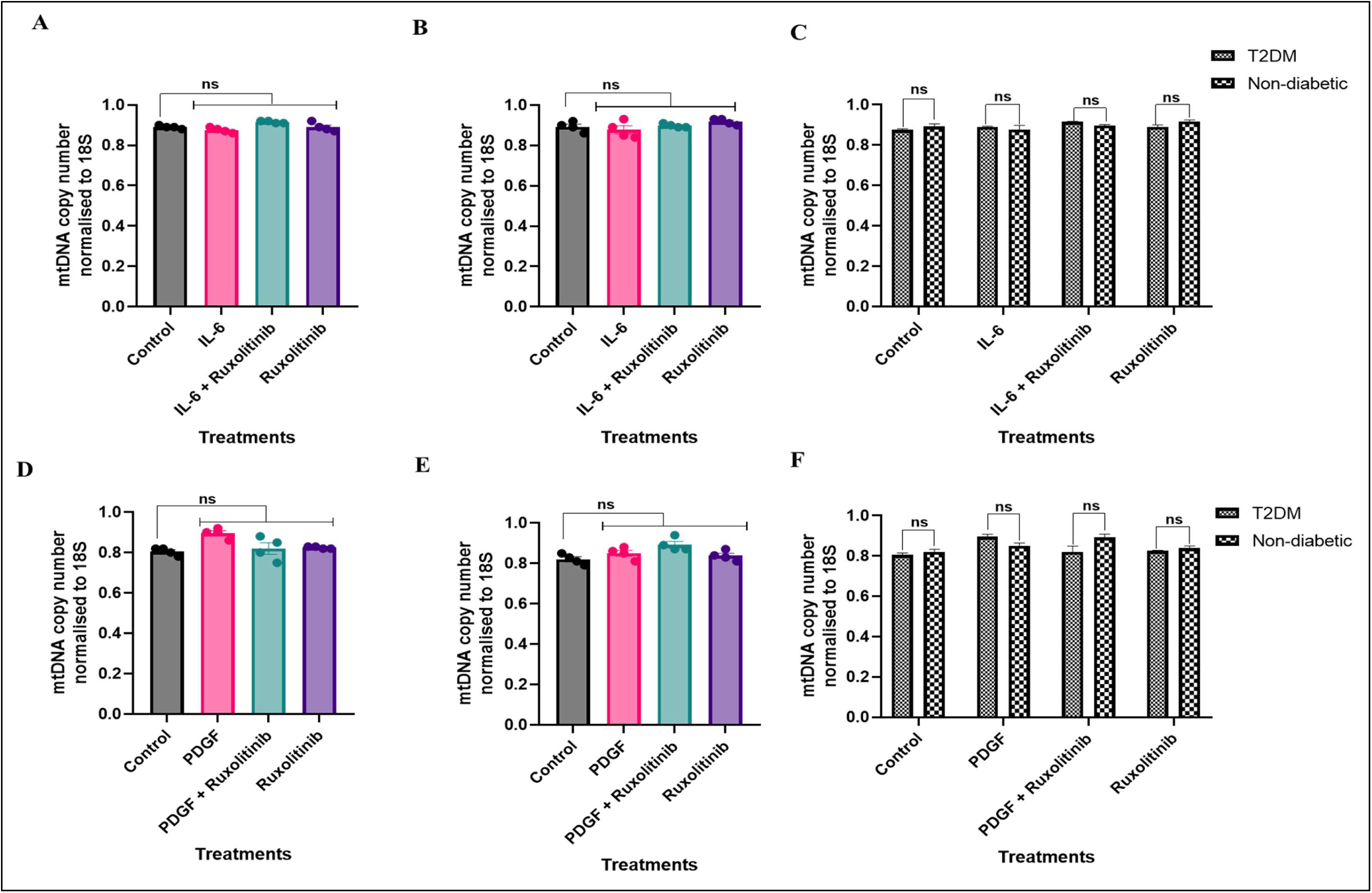
mtDNA copy number of HSVSMCs from T2DM and non-diabetic patients after treatment with IL-6/sIL-6Rα orPDGF-BB +/- ruxolitinib. qPCR analysis of mtDNA copy number of mitochondrially encoded gene COI as a marker of mtDNA copy number normalised to 18S. **Top panel:** (A) T2DM, (B) non-diabetic patients, and (C) comparison between T2DM patients versus non-diabetic control treated with IL-6/sIL-6Rα (IL-6 (5 ng/ml) and sIL-6Rα (25 ng/ml)) +/- ruxolitinib (0.1 μM) **Bottom panel:** (D) T2DM, (E) non-diabetic patients, and (F) comparison between T2DM patients versus non-diabetic control treated with PDGF-BB (10 ng/ml) +/- ruxolitinib (0.1 μM). All data were normalised to 18S, and normalised data from different treatment conditions were compared with data from normalised untreated control for statistical difference. Data are presented as mean ± SEM from n=4 experiments using HSVSMC from different patients. Statistical significance was assessed using one-way ANOVA followed by Dunnett’s post-hoc test to determine significant differences between means IL-6: IL-6/sIL-6Rα; PDGF: PDGF-BB.

### 3.6. Effect of Ang ll on OCR of HSVSMCs from T2DM and non-diabetic patients

While Ang II has been shown to promote VSMC migration and proliferation by activating the MAP/ERK1,2 pathway, ^37^ it is not totally clear how Ang II modulates these processes. However, we hypothesise that the increased cell proliferation and migration induced by Ang II ^37^ may necessitate higher ATP synthesis, which would increase oxygen consumption. Therefore, we assessed the effect of Ang II +/- MEK1/2-selective inhibitor trametinib on the OCR of HSVSMCs from patients with and without T2DM. The optimal concentration of trametinib that significantly (*p*<0.05 versus untreated cells, n=4) inhibited activation of ERK1/2 in HSVSMCs from non-diabetic patients was 10 nM trametinib (Supplementary Information Figure S2). Hence, this concentration was used in subsequent experiments. As shown in Figures 6A and B, Ang II (100 nM) did not significantly alter the OCR of HSVSMCs from T2DM and non-diabetic patients, respectively, even though the same concentration of Ang II stimulated a transient increase in ERK1/2 phosphorylation in HSVSMCs (Supplementary Information Figure S3). However, trametinib with or without Ang II stimulation significantly reduced OCRs at basal respiration (*p*<0.01 versus unstimulated and Ang II-stimulated cells, n=4) and maximal respiration (*p*<0.05 versus unstimulated and Ang II-stimulated cells, n=4) in HSVSMCs from T2DM (Figure 6A). Similarly, in HSVSMC from non-diabetic patients, Ang II treatment did not significantly alter the OCR (Figure 6B). Furthermore, trametinib caused a significant reduction in the OCRs at basal respiration (*p*<0.01 with Ang II stimulation; *p*< 0.001 without Ang II stimulation, both versus unstimulated and Ang II-stimulated cells, n=4) in HSVSMCs from non-diabetic patients (Figure 6B). Interestingly, trametinib caused a significant reduction in the OCRs at maximal respiration (*p*<0.05 with or without Ang II stimulation versus unstimulated and Ang II-stimulated cells, n=4) in HSVSMCs from non-diabetic patients (Figure 6B). Also, direct comparison of OCRs after treatment with Ang II revealed a significant increase in the OCR of HSVSMCs from T2DM at maximal respiration (*p*<0.05 versus HSVSMCs from non-diabetic patients, n=4; Figure 6C).

**Figure 6:**
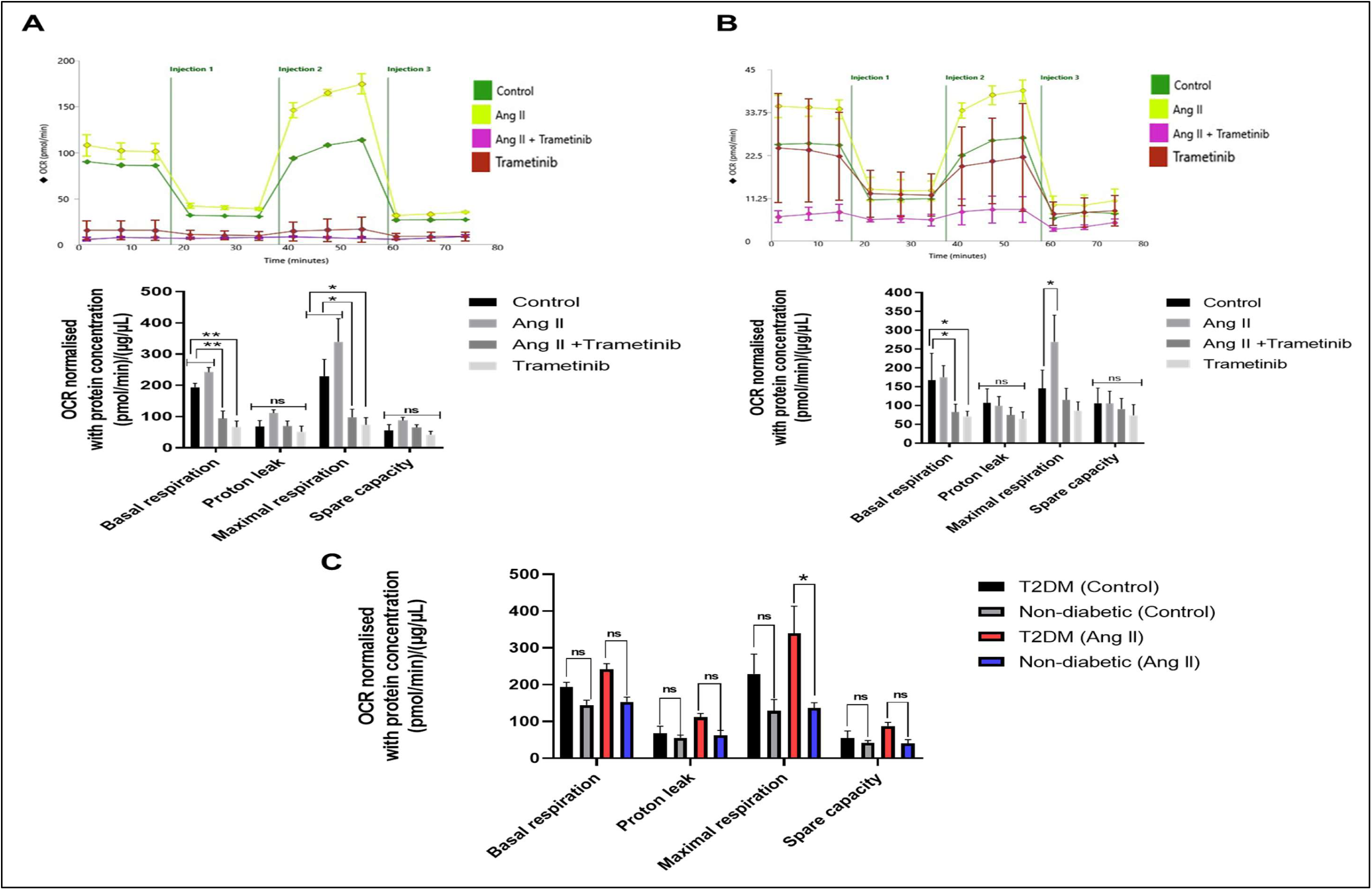
Mitochondria stress analysis to determine the OCR of HSVSMCs from T2DM and non-T2DM patients after stimulation with Ang II +/- trametinib. **(M) Upper panel:** representative time course curve of OCR of HSVSMCs from T2DM patients treated with Ang II (100 nM) +/- trametinib (10 nM) and untreated control. **Lower panel:** comparative analysis of normalised peak OCR between treatment groups and untreated control at basal glycolysis, and after addition of inhibitors of mitochondrial respiration and the uncoupler. Normalised data are presented as mean ± SEM from n=4 biological replicates using HSVSMC samples from different T2DM patients. **(N) Upper panel:** representative time course curve of OCR of HSVSMCs from non-diabetic patients treated with Ang II (100 nM) +/- trametinib (10 nM) and untreated control. **Lower panel:** comparative analysis of normalised peak OCR between treatment groups and untreated control at basal glycolysis, and after addition of inhibitors of mitochondrial respiration and the uncoupler. Normalised data are presented as mean ± SEM from n=4 biological replicates using HSVSMC samples from different non-diabetic patients. **(O)** Comparison of normalised peak OCR of unstimulated and Ang II-stimulated HSVSMCs from T2DM patients versus non-diabetic control. Normalised data are presented as mean ± SEM and statistical significance was assessed using an independent t-test, n=4 biological replicates using HSVSMC samples from different T2DM and non-diabetic patients. In the lower panels of Figures A and B, normalised data from different treatment conditions were compared with data from the normalised untreated control. Statistical significance was assessed using one-way ANOVA followed by Dunnett’s post-hoc test to determine significant differences between means, **\****P* < 0.05; \*\**P* < 0.01. Ang II: Angiotensin II; OCR: oxygen consumption rate.

### 3.7. Effect of Ang ll on ECAR of HSVSMCs from T2DM and non-diabetic patients

There is evidence that Ang II may cause mitochondrial oxidative damage in vascular endothelial cells, which could reduce NO bioavailability and increase vascular oxidative stress.

^21^ While this finding emphasises the role of Ang II in controlling metabolic homeostasis in endothelial cells, its effect on ECAR of HSVSMCs is unknown. Therefore, we evaluated the effects of Ang ll +/- trametinib on ECAR of HSVSMCs from T2DM and non-diabetic control. As shown in Figure 7A, Ang II treatment did not significantly alter the ECAR in HSVSMCs from T2DM patients. However, in HSVSMCs from T2DM, trametinib significantly reduced ECARs at basal glycolysis (*p*<0.05 with Ang II stimulation, *p*<0.01 without Ang II stimulation, n=4), and glycolytic capacity (*p*<0.05 with Ang II stimulation, *p*<0.01 without Ang II stimulation, n=4). Furthermore, trametinib significantly reduced ECARs at maximal glycolysis (*p*<0.05 with Ang II stimulation, *p*<0.01 without Ang II stimulation, n=4), and glycolytic reserve (*p*<0.05 with Ang II stimulation, *p*<0.01 without Ang II stimulation, n=4) in HSVSMCs from T2DM. Similarly, in HSVSMCs from non-diabetic patients, there was no significant change to ECAR in response to Ang II (Figure 7B). Conversely, trametinib caused a significant reduction in the ECARs at basal glycolysis (*p*<0.01 with or without Ang II stimulation, n=4), glycolytic capacity (*p*<0.01 with or without Ang II stimulation, n=4), maximal glycolysis (*p*<0.05 with or without Ang II stimulation, n=4), and glycolytic reserve (*p*<0.05 with or without Ang II stimulation, n=4) in HSVSMCs from non-diabetic patients (Figure 7B). However, no significant differences were observed in the ECARs between Ang II-stimulated HSVSMCs from T2DM and non-diabetic patients (Figure 7C).

**Figure 7:**
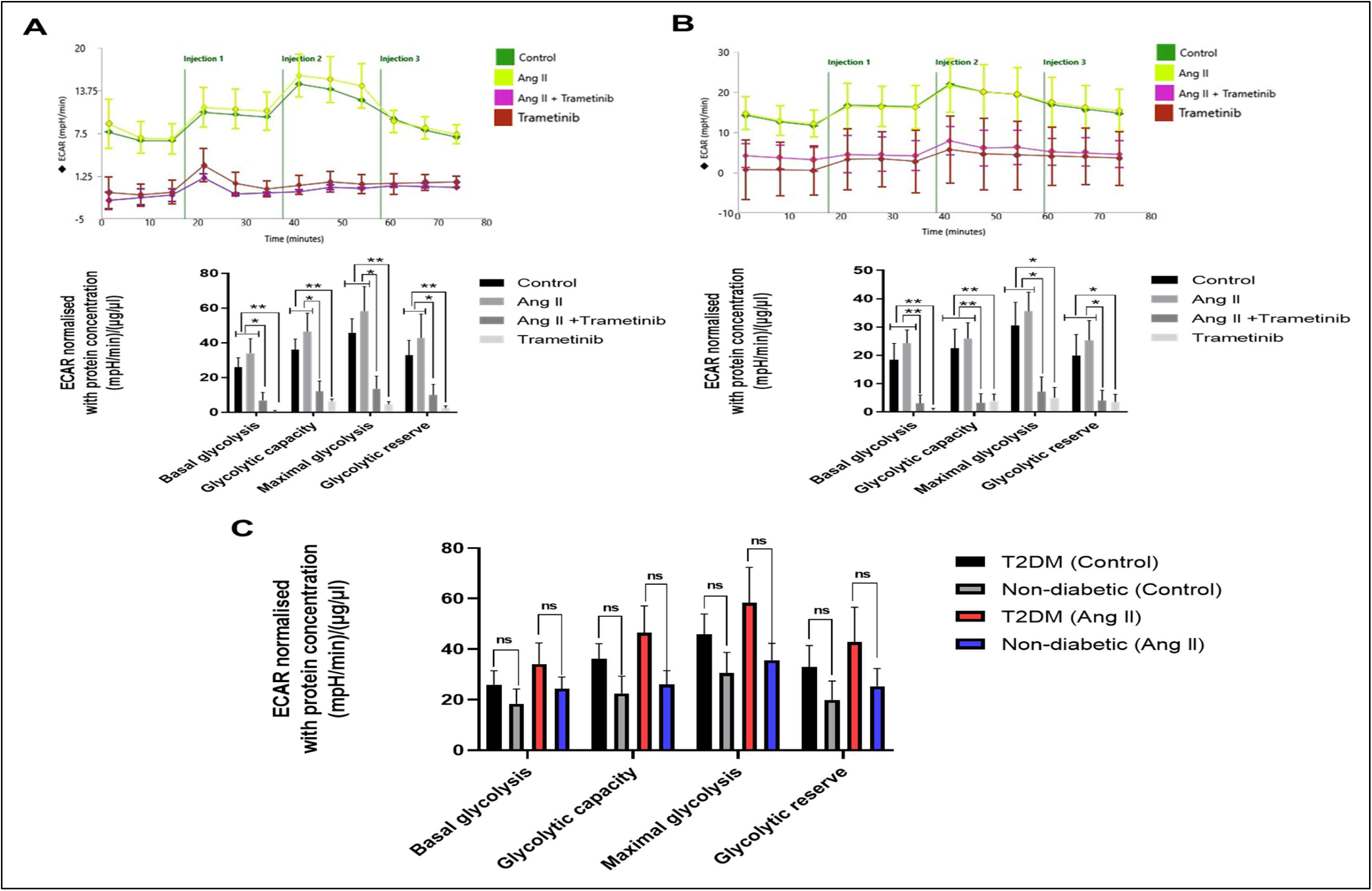
Mitochondria stress analysis to determine the ECAR of HSVSMCs from T2DM and non-T2DM patients after stimulation with Ang II +/- trametinib. **(P) Upper panel:** representative time course curve of ECAR of HSVSMCs from T2DM patients treated with Ang II (100 nM) +/- trametinib (10 nM) and untreated control. **Lower panel:** comparative analysis of normalised peak ECAR between treatment groups and untreated control at basal glycolysis, and after addition of inhibitors of mitochondrial respiration and the uncoupler. Normalised data are presented as mean ± SEM from n=4 biological replicates using HSVSMC samples from different T2DM patients. **(Q) Upper panel:** representative time course curve of ECAR of HSVSMCs from non-diabetic patients treated with Ang II (100 nM) +/- trametinib (10 nM) and untreated control. **Lower panel:** comparative analysis of normalised peak ECAR between treatment groups and untreated control at basal glycolysis, and after addition of inhibitors of mitochondrial respiration and the uncoupler. Normalised data are presented as mean ± SEM from n=4 biological replicates using HSVSMC samples from different non-diabetic patients. **(R)** Comparison of normalised peak ECAR of unstimulated and PDGF-BB-stimulated HSVSMCs from T2DM patients versus non-diabetic control. Normalised data are presented as mean ± SEM and statistical significance was assessed using an independent t-test, n=4 biological replicates using HSVSMC samples from different T2DM and non-diabetic patients. In the lower panels of Figures A and B, normalised data from different treatment conditions were compared with data from the normalised untreated control. Statistical significance was assessed using one-way ANOVA followed by Dunnett’s post-hoc test to determine significant differences between means, **\****P* < 0.05; \*\**P* < 0.01. Ang II: Angiotensin II; ECAR: extracellular acidification rate.

### 3.8. Effect of thrombin on OCR of HSVSMCs from T2DM and non-diabetic patients

Thrombin has been associated with vascular remodelling via a number of potential mechanisms, including activation of the MAPK/ERK signalling pathway ^25^. However, it is still unknown how thrombin affects mitochondrial respiration, a crucial mediator of VSMC metabolic function. Given that understanding the metabolic homeostasis of VSMCs is crucial for targeting vascular remodelling, we therefore assessed the impact of thrombin +/- trametinib on the OCR of HSVSMCs from T2DM and non-diabetic controls. Thrombin caused a significant increase (p<0.05 versus unstimulated cells) in OCR of HSVSMCs from T2DM patients at maximal respiration which was abolished by trametinib (Figure 8A). On the other hand, trametinib significantly reduced OCR at basal respiration (*p*<0.05 with or without thrombin stimulation versus unstimulated cells, n=4). Furthermore, in HSVSMCs from non-diabetic patients, thrombin did not significantly change OCR, although trametinib significantly reduced (at both basal and maximal respiration OCRs (*p*<0.05 with or without thrombin stimulation versus unstimulated cells, n=4; Figure 8B). Meanwhile, direct comparison of OCRs in thrombin-treated HSVSMCs from T2DM and non-diabetic patients revealed there was a significant increase in OCR at maximal respiration (*p*<0.01, n=4), and spare capacity (*p*<0.05, n=4) (Figure 8C). Moreover, spare capacity was significantly higher in control T2DM HSVSMCs versus non-diabetic control cells.

**Figure 8:**
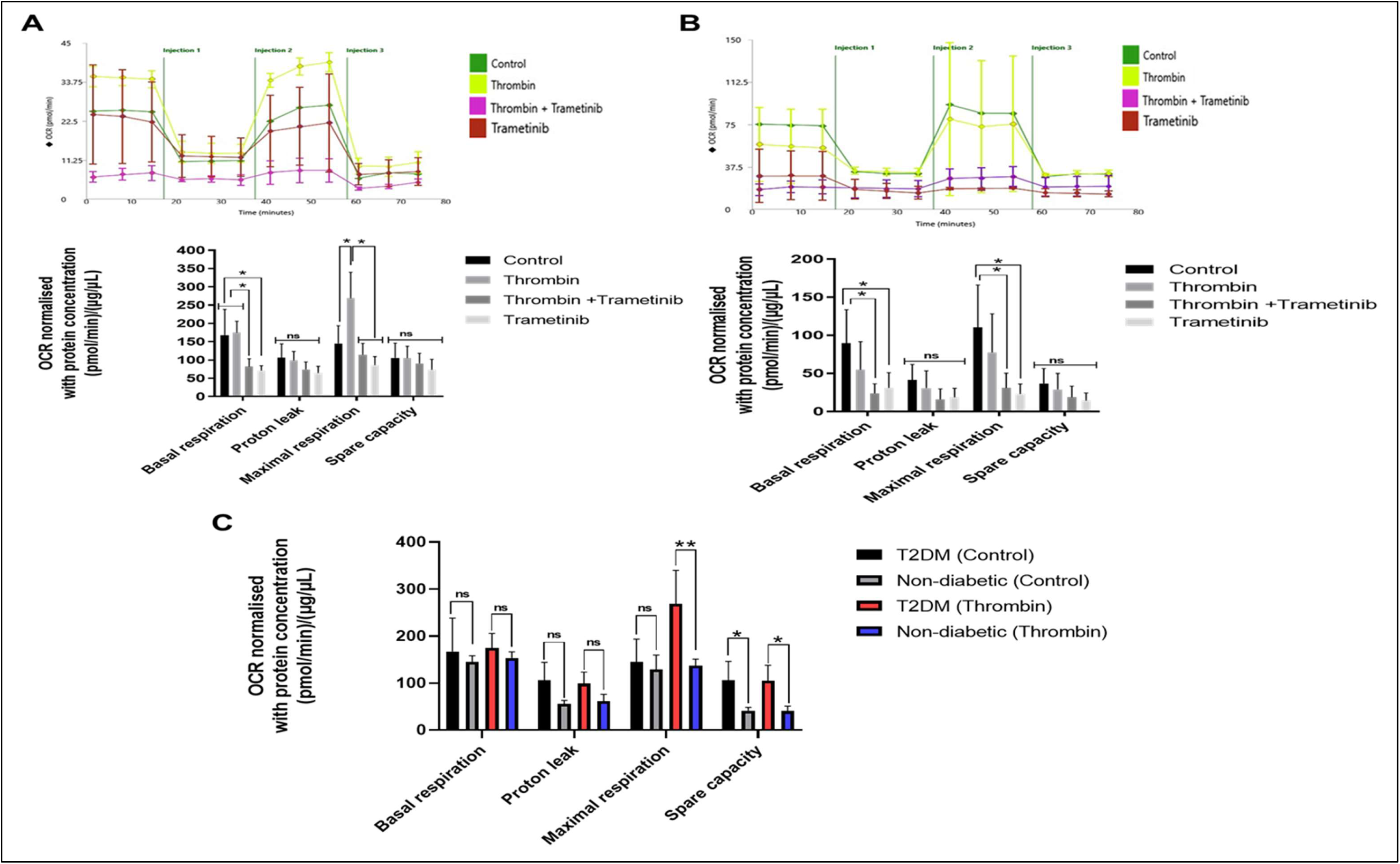
Mitochondria stress analysis to determine the OCR of HSVSMCs from T2DM and non-T2DM patients after stimulation with thrombin +/- trametinib. **(S) Upper panel:** representative time course curve of OCR of HSVSMCs from T2DM patients treated with thrombin (1 U/ml) +/- trametinib (10 nM) and untreated control. **Lower panel:** comparative analysis of normalised peak OCR between treatment groups and untreated control at basal glycolysis, and after addition of inhibitors of mitochondrial respiration and the uncoupler. Normalised data are presented as mean ± SEM from n=4 biological replicates using HSVSMC samples from different T2DM patients. **(T) Upper panel:** representative time course curve of OCR of HSVSMCs from non-diabetic patients treated with thrombin (1 U/ml) +/- trametinib (10 nM) and untreated control. **Lower panel:** comparative analysis of normalised peak OCR between treatment groups and untreated control at basal glycolysis, and after addition of inhibitors of mitochondrial respiration and the uncoupler. Normalised data are presented as mean ± SEM from n=4 biological replicates using HSVSMC samples from different non-diabetic patients. **(U)** Comparison of normalised peak OCR of unstimulated and thrombin-stimulated HSVSMCs from T2DM patients versus non-diabetic control. Normalised data are presented as mean ± SEM and statistical significance was assessed using an independent t-test, n=4 biological replicates using HSVSMC samples from different T2DM and non-diabetic patients. In the lower panels of Figures A and B, normalised data from different treatment conditions were compared with data from the normalised untreated control. Statistical significance was assessed using one-way ANOVA followed by Dunnett’s post-hoc test to determine significant differences between means, **\****P* < 0.05. ECAR: extracellular acidification rate.

### 3.9. Effect of thrombin on ECAR of HSVSMCs from T2DM and non-diabetic patients

Currently, there are no documented research findings highlighting the effects of the MAPK/ERK pathway on ECAR of VSMCs despite the significance of ECAR as a tool for measuring glycolytic flux in cells. ^38^ Therefore, we assessed the effect of thrombin +/- trametinib on the ECAR of HSVSMCs from T2DM and non-diabetic controls. As shown in Figure 9A, thrombin did not cause any significant alteration in the ECAR of HSVSMCs from T2DM patients. However, trametinib significantly reduced ECARs at basal glycolysis, glycolytic capacity, maximal glycolysis, and glycolytic reserve (*p*<0.01 with or without thrombin, n=4) in HSVSMCs from T2DM patients (Figure 9A). Similarly, in HSVSMCs from non-diabetic patients, while there was no significant alteration in ECAR after thrombin stimulation, trametinib significantly reduced the OCRs at basal glycolysis (*p*<0.01 with or without thrombin, n=4), maximal glycolysis (*p*<0.05 with or without thrombin, n=4), and glycolytic reserve (*p*<0.05 with thrombin, n=4), in HSVSMCs from non-diabetic patients (Figure 9B). Also, the ECAR in thrombin-stimulated HSVSMCs from T2DM patients was significantly higher at maximal glycolysis when compared with those from non-diabetic patients (*p*<0.05, n=4; Figure 9C).

**Figure 9:**
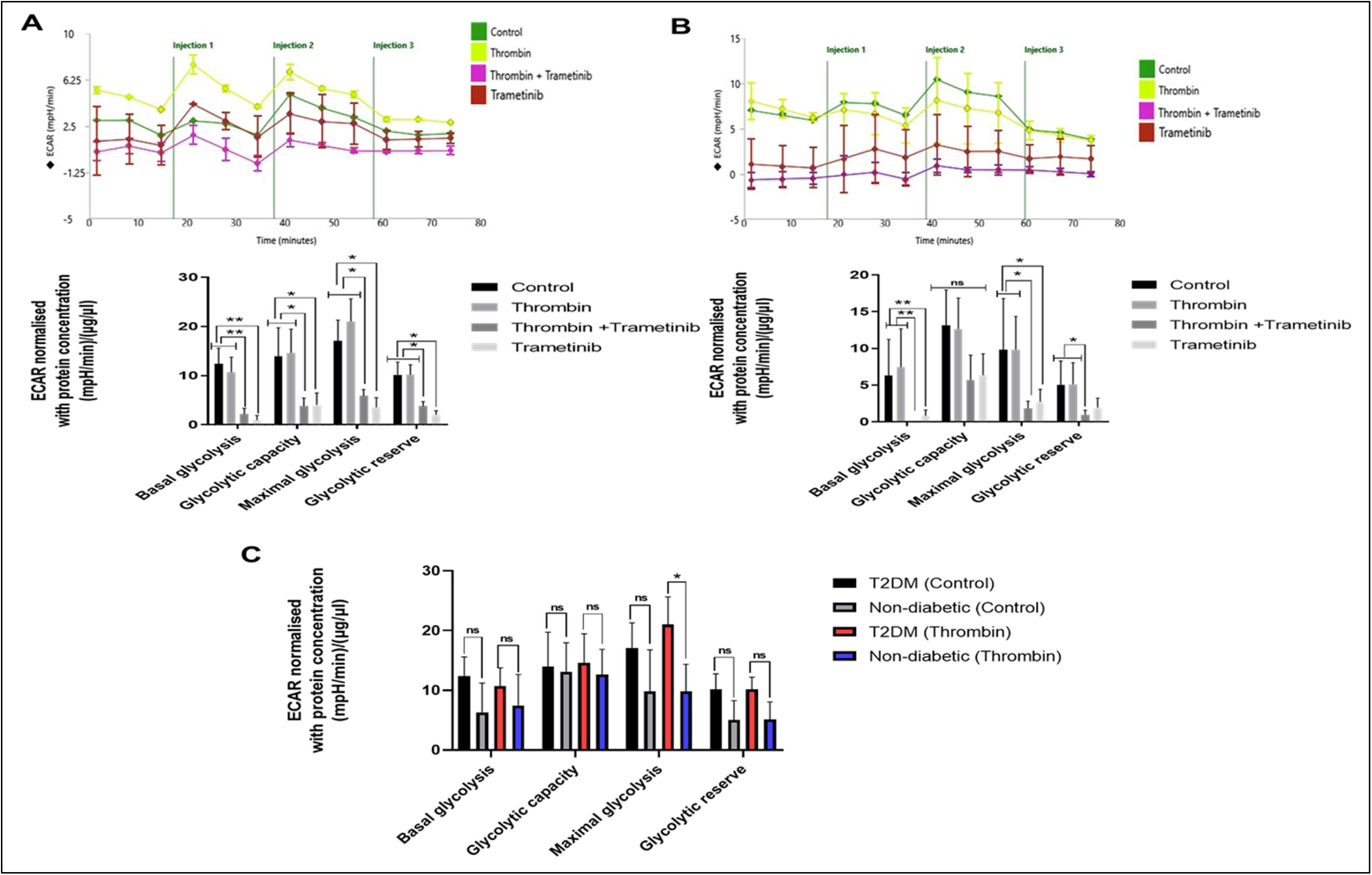
Mitochondria stress analysis to determine the ECAR of HSVSMCs from T2DM and non-T2DM patients after stimulation with thrombin +/- trametinib. **(V) Upper panel:** representative time course curve of ECAR of HSVSMCs from T2DM patients treated with thrombin (1 U/ml) +/- trametinib (10 nM) and untreated control. **Lower panel:** comparative analysis of normalised peak ECAR between treatment groups and untreated control at basal glycolysis, and after addition of inhibitors of mitochondrial respiration and the uncoupler. Normalised data are presented as mean ± SEM from n=4 biological replicates using HSVSMC samples from different T2DM patients. **(W) Upper panel:** representative time course curve of ECAR of HSVSMCs from non-diabetic patients treated with thrombin (1 U/ml) +/- trametinib (10 nM) and untreated control. **Lower panel:** comparative analysis of normalised peak ECAR between treatment groups and untreated control at basal glycolysis, and after addition of inhibitors of mitochondrial respiration and the uncoupler. Normalised data are presented as mean ± SEM from n=4 biological replicates using HSVSMC samples from different non-diabetic patients. **(X)** Comparison of normalised peak ECAR of unstimulated and thrombin-stimulated HSVSMCs from T2DM patients versus non-diabetic control. Normalised data are presented as mean ± SEM and statistical significance was assessed using an independent t-test, n=4 biological replicates using HSVSMC samples from different T2DM and non-diabetic patients. In the lower panels of Figures A and B, normalised data from different treatment conditions were compared with data from the normalised untreated control. Statistical significance was assessed using one-way ANOVA followed by Dunnett’s post-hoc test to determine significant differences between means, **\****P* < 0.05; \*\**P* < 0.01. ECAR: extracellular acidification rate.

### 3.10. mtDNA copy number in HSVSMCs from T2DM and non-diabetic patients after treatments with Ang II and thrombin +/-trametinib

Ang II caused a significant increase in the OCR of HSVSMCs from T2DM at maximal respiration (p<0.05 versus HSVSMCs from non-diabetic patients, n=4; Figure 6C). Also, the OCR of HSVSMCs from T2DM, but not from non-diabetic, was significantly increased by thrombin (Figure 8). It was therefore important to determine whether this could be explained by an increase in mitochondria number in response to stimulation by Ang II and thrombin. Therefore, the copy number of the mitochondrially-encoded gene COI after treatment with Ang II and thrombin +/-trametinib was determined using qPCR. ^35, 36^

As shown in Figure 10A, there was no significant difference in the mtDNA copy number in HSVSMCs from T2DM patients after treatments with Ang II+/-trametinib compared with untreated control. Also, in the HSVSMCs from non-diabetic patients, there was no significant difference in mtDNA copy number after treatment with Ang II+/-trametinib compared with untreated control (Figure 10B). Comparison between HSVSMCs from T2DM patients versus non-diabetic control treated with Ang II +/- trametinib showed no significant difference (Figure 10C). As shown in Figure 10D, there was no significant difference in mtDNA copy number in HSVSMCs from T2DM patients after treatments with thrombin+/-trametinib compared with untreated control. There was also no significant difference in the mtDNA copy number in HSVSMCs from non-diabetic patients after treatment with thrombin+/-trametinib compared with untreated control (Figure 10E). Also, comparison between HSVSMCs from T2DM patients versus non-diabetic control treated with thrombin +/- trametinib showed no significant difference (Figure 10F).

**Figure 10:**
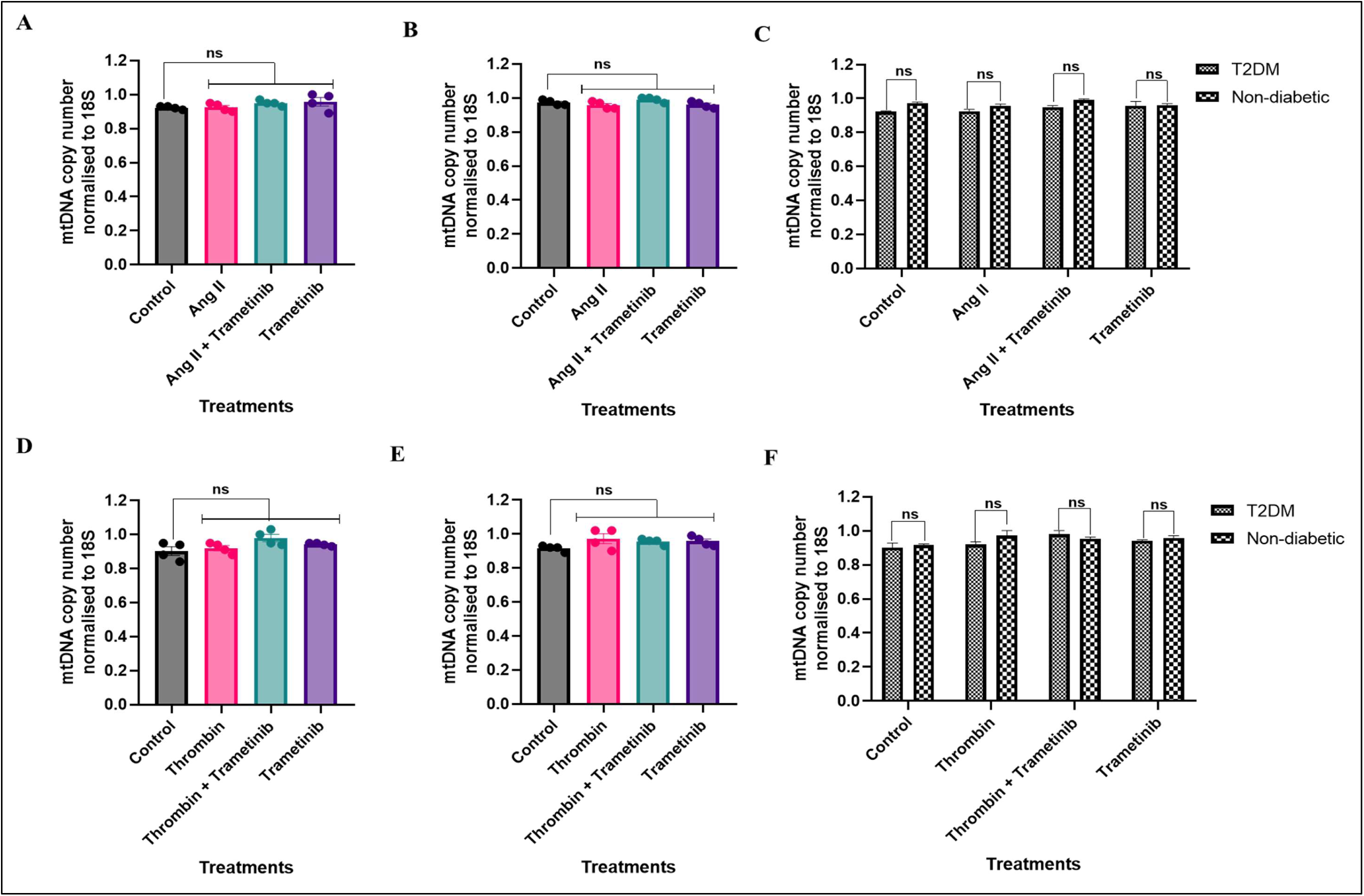
mtDNA copy number in HSVSMCs from T2DM and non-diabetic patients after treatment with Ang II and thrombin +/- trametinib. mtDNA copy number of mitochondrially encoded gene COI as a marker of mtDNA copy number in **Top panel: (A)** T2DM, and **(B)** non-diabetic patients, and (C) comparison between T2DM patients versus non-diabetic control treated with Ang II (100 nM) +/- trametinib (10 nM). **Bottom Panel: (**D) T2DM, **(E)** non-diabetic patients, **(F)** comparison between T2DM patients versus non-diabetic control treated with thrombin (1 U/ml) +/- trametinib (10 nM). All data were normalised to 18S, and normalised data from different treatment conditions were compared with data from normalised untreated control for statistical difference. Data are presented as mean± SEM from n=4 experiments using HSVSMC from different patients. Data are presented as mean± SEM from n=4 experiments using HSVSMC from different patients. Statistical significance was assessed using one-way ANOVA followed by Dunnett’s post-hoc test to determine significant differences between means. Ang II, Angiotensin II.

## 4. Discussion

VSMCs have garnered attention as a possible target to treat vascular pathologies due to their role in neointima hyperplasia (NIH) and venous graft stenosis. ^39–42^ More so, HSV wall thickening is caused by the HSVSMCs shifting from a differentiated to a dedifferentiated phenotype, resulting in uncontrolled cell migration and proliferation responsible for long-term VGF. ^43^ The rate of VGF after 10 years is about 50% and is suggested to be worse (70%) in T2DM patients. ^44, 45^ Furthermore, it has been established that VSMCs from T2DM patients exhibit unique functional characteristics and are more migratory than those of non-diabetic controls. ^6, 7^ Additionally, there is evidence that T2DM trigger changes in mitochondrial function of VSMCs of mice. ^46^ However, for HSVSMCs, an important type of cell linked to the vascular dysfunction causing saphenous VGF, the effects of T2DM on the mitochondrial function and potential downstream signalling mechanisms that regulate these processes are unknown. Hence, our findings have highlighted some of the downstream signalling pathways that are involved in regulating these mitochondrial functions alongside possible effects of T2DM on these functions.

While it is well established that the levels and activities of proinflammatory cytokines such as IL-6 are increased in T2DM, ^47^ the exact impact on mitochondrial OCR and ECAR, in VSMCs and specifically HSVSMCs is unclear. Our study has now shown that IL-6/sIL-6Rα increases mitochondrial respiration through the JAK/STAT pathway, as the IL-6/sIL-6Rα-induced increase in OCR of HSVSMCs from T2DM but not in non-diabetic patients was abolished by JAK1/2-selective inhibitor ruxolitinib (Figure 1). Ruxolitinib also caused a reduction of basal OCR of HSVSMCs from both T2DM and non-diabetic patients (Figure 1). The JAK/STAT pathway is an emerging target in inflammation, a major feature of cardiovascular diseases. Hence, anti-inflammatory drugs are increasingly being considered in the management of cardiovascular events including those resulting from vascular dysfunction. ^48^ For instance, current available treatments for pulmonary hypertension (PH), which is caused by remodeling of the pulmonary arteries marked by endothelial dysfunction and proliferation of smooth muscle cells, are vasodilatory drugs, which do not stop disease progression. ^49^ It has been well demonstrated that the JAK/STAT molecular pathway plays an essential role in the vascular remodelling linked to pulmonary hypertension. ^49^ Interestingly, our results showed that stimulation of HSVSMCs with IL-6/sIL-6Rα did not significantly modify the ECAR of HSVSMCs from either T2DM and non-diabetic patients, suggesting that glycolysis is unaffected by the JAK/STAT pathway. The reason for this is still unclear, however, it has been shown that IL-6/sIL-6Rα shifted the control for ATP synthesis towards processes that generate the mitochondrial membrane potential. ^50^ This suggests that IL-6/sIL-6Rα induces a metabolic state where cellular functions are limited by the mitochondrial energy supply, limiting the conversion of glucose to lactate, which determines ECAR. ^50^

Similarly, PDGF-BB increased OCR of HSVSMCs from T2DM at basal and maximal respiration and these were again both abolished by ruxolitinib (Figure 3), which suggests that the modulation of mitochondrial respiration by PDGF-BB is also JAK 1/2-dependent. Conversely, there was no significant increase in the OCR of HSVSMCs from non-diabetic subjects in response to PDGF-BB (Figure 3). Also, in both HSVSMCs from T2DM and non-diabetic patients, ruxolitinib caused a significant reduction in the basal and PDGF-stimulated OCR under all conditions (Figure 3). The importance of mitochondrial metabolism as a key regulator of cell growth and proliferation is becoming increasingly appreciated. ^51^ Increased glycolysis and glutamine usage are seen in rapidly dividing cells, which contribute to the production of energy, reducing equivalents, and the carbon and nitrogen building blocks necessary for daughter cell synthesis. ^51^ Growth factors like PDGF have been found to boost glycolysis and mitochondrial activity in rat aortic VSMCs. ^52^ Additionally, studies on pre-clinical models and human pulmonary arterial SMCs have demonstrated that VSMCs have greater rates of glycolysis. ^53, 54^ These studies also indicated a role for mitochondrial fission in pulmonary arterial SMC proliferation, with pulmonary arterial SMC mitochondria in cellular and pre-clinical pulmonary arterial hypertension (PAH) models found to be dysmorphic and hyperpolarised. ^53, 54^ While little is known about how mitochondrial structure controls the phenotypic and metabolic properties of HSVSMCs, it has been demonstrated that such changes in mitochondrial morphology have a direct impact on cell differentiation and proliferation of rat aortic SMC. ^55, 56^ Furthermore, despite numerous studies that have linked PDGF to vascular dysfunction, ^56–60^ a role for PDGF on mitochondrial respiration and its metabolic components has not yet been investigated in HSVSMCs. The significance of our findings is highlighted by the discovery that PDGF-BB can control mitochondrial respiration in HSVSMCs via the JAK/STAT pathway, which is less well-known for PDGF’s downstream activity than other pathways such as ERK1,2 and Akt. ^61^

Conversely, ruxolitinib reduced ECAR in HSVSMCs from both T2DM and non-diabetic patients (Figures 2 and 3) and also abolished the PDGF-BB-induced increase of ECAR in HSVSMCs from T2DM patients (Figure 3). These findings suggest a JAK-mediated modulation of mitochondrial glycolysis of HSVSMCs. The JAK/STAT pathway is an established therapeutic target for immune and inflammatory conditions that promote several CVDs. ^48^ Furthermore, emerging findings have suggested that patients with PAH of different types have pulmonary arteries with overactive JAK/STAT pathways. ^49, 62^ Additionally, several cytokines, including IL-6, IL-13, and IL-11, as well as growth factors, including PDGF, VEGF, and TGFβ1, activate the JAK/STAT pathway and trigger pulmonary remodelling, which contributes to the onset of PAH. ^49^ Therefore, targeting the cellular signalling pathways implicated in vascular dysfunction to prevent vascular remodelling may also be a viable option for preventing VGF.

Furthermore, our results showed that Ang II did not cause any noticeable alteration in OCR of HSVSMCs from both T2DM and non-diabetic patients. However, trametinib reduced OCR at basal and maximal respiration of HSVSMCs from both T2DM and non-diabetic patients (Figure 6). This suggests that mitochondrial respiration can be modulated through the ERK1,2 signalling pathway. Furthermore, OCR of HSVSMCs from T2DM and non-diabetic patients did not significantly differ from each other in either the absence or presence of Ang II (Figure 6). While we have described this in HSVSMCs for the first time, it was previously demonstrated that Ang II caused a reduction in the OCR of HUVECs. ^63^ In this study, ^65^ it was shown that Ang II reduced OCR at basal and maximal respiration compared with the control group. The explanation for this is unclear at this time.

In addition, the ERK1,2 pathway can indirectly modulate cellular metabolic homeostasis through tumor-suppressor liver kinase B1-induced activation of AMPK, a crucial regulator of energy homeostasis, under low-energy situations in most cellular contexts. ^64,65^ When activated, AMPK blocks most of the biosynthetic pathways required for cell growth, reducing ATP utilisation, and activates catabolic pathways that produce ATP, enabling cells to restore energy equilibrium. ^65^ Furthermore, it has been shown that Ang II decreased OCR of mitochondria isolated from the kidney cortex of both non-diabetic controls and type 1 diabetic rats via AT2 receptor-mediated nitric oxide release. ^66^ It was further demonstrated in this study that whereas Ang II increases OCR at mitochondria leak respiration in diabetic animals via AT1 receptors, AT1 receptors do not influence mitochondria function in control mice. ^66^ Together, these findings suggest that while Ang II can elicit its effects on mitochondrial function activity through multiple signalling pathways, in the current study we have looked specifically at the effect of Ang II as a known activator of the ERK1,2 signalling in HSVSMCs, where it did not cause any significant alteration in the OCR in cells from T2DM or non-diabetic patients. Also, while there were no significant alterations in ECAR of HSVSMCs from both T2DM and non-diabetic patients after stimulation with Ang II, trametinib caused a significant reduction in ECAR in HSVSMCs from both T2DM and non-diabetic patients (Figure 7).

We also found that thrombin significantly increased OCR in HSVSMCs from T2DM but not non-diabetic patients, and that this was attenuated by trametinib (Figure 10). Direct comparison of the OCR of HSVSMCs from T2DM and non-diabetic patients revealed that thrombin significantly increased the OCR of HSVSMCs from T2DM patients compared to non-diabetic control subjects at maximal glycolysis and spare capacity (Figure 8). In addition, trametinib reduced basal OCR in HSVSMCs from both T2DM and non-diabetic patients (Figure 8). Previous findings have revealed that thrombin triggered a significant rise in OCR in platelets. ^67, 68^ While these two studies used platelets from healthy donors, ^67, 68^ similar studies using unstimulated platelets from T2DM patients indicated reduced OCR when compared with those from healthy donors. ^69^ We have now demonstrated an increase in the OCR of HSVSMCs from T2DM patients, but not non-diabetic patients, following thrombin stimulation. The fact that trametinib attenuated this increase in OCR suggests that thrombin elicited its effect through the ERK1,2 pathway, although the underlying mechanism is not entirely clear.

Trametinib, the MEK1/2 inhibitor employed in this study not only abolished the thrombin-induced increase in OCR of HSVSMCs from T2DM but also reduced the OCR of HSVSMCs from both T2DM and non-diabetic patients. Trametinib binds to unphosphorylated MEK1/2 preferentially, blocking Raf-dependent phosphorylation and activation of MEK. ^70^ While it might be premature to assume that the ability of trametinib to abolish the MEK-mediated increase in OCR in HSVSMCs from T2DM patients would be of any clinical benefit, however, trametinib used alone or in combination with B-Raf inhibitor dabrafenib is a clinical tool used to manage malignant melanoma driven by constitutively active Val600Glu- and Val600Lys-mutated B-Raf. ^71^ Dabrafenib binds the ATP binding site of the mutated B-Raf protein, specifically the Val600Glu mutation. This inhibits the kinase activity of B-Raf and ultimately ERK1/2, resulting in programmed cell death. ^72^

It has long been recognised that mitochondrial dysfunction contributes significantly to the underlying pathogenesis of a number of age-related diseases, including cancers, CVDs and neurodegenerative diseases. ^73, 74^ Hence, mtDNA copy number is increasingly used to evaluate the role of mitochondria in diseases as its measurement is a straightforward proxy for mitochondrial function. ^73^ Additionally, mtDNA copy number provides an opportunity to determine changes not only in disease conditions but as a genetic marker. In this study, we used the mtDNA copy number to investigate if the observed alterations in OCR and ECAR detected in HSVSMCs from T2DM after the activation of the JAK/STAT and the MAPK/ERK signalling pathways was caused by an increase in mtDNA copy number or by other factors that are unknown but may be related to T2DM. From our findings, there were no significant changes in the mtDNA copy number after treatment with the different treatment conditions, and when compared between T2DM patients and non-T2DM controls (Figures 5 and 10). These data therefore confirm that the observed changes in OCR relate to increased metabolic activity by the mitochondria. Therefore, while it is not entirely clear, the observed increase in OCR and ECAR in HSVSMCs from T2DM and not from non-diabetic control could be due to the T2DM status.

In summary, understanding any link between T2DM and compromised vascular function, specifically in autologous saphenous veins used as conduits in bypass surgery procedures in T2DM patients, is critical to improving vein graft patency and patient outcomes. Our study has identified downstream signalling pathways involved in regulating mitochondrial function in HSVSMCs alongside described how T2DM potentially modifies these functions. The significance of these findings lies in the fact that proliferation and migration of vascular cells including SMCs have been linked to vascular remodelling and restenosis responsible for VGF. Furthermore, SMCs from T2DM have been demonstrated to be more migratory when compared to those from non-diabetic patients. ^6, 7^ Putting these together, as more oxygen and enhanced glycolysis is required to generate more ATP to drive these cellular processes, identifying the pathways where this can be effectively modulated is of significance in the development of novel therapeutic opportunities to limit vein graft failure.

## Conclusion

Through this study, we have demonstrated a JAK- and MEK-mediated regulation of mitochondrial function (OCR and ECAR) alongside possible effect of T2DM on these processes in HSVSMC, a key cell type implicated in vascular dysfunction that is responsible for VGF. Our findings have therefore contributed to the body of knowledge identifying possible novel glucose-dependent alterations that cause vascular dysfunction in T2DM patients. These findings therefore offer a background and rationale for future investigations to develop novel therapies to improve vein graft patency.

## Abbreviations

AMPK: adenosine monophosphate-activated protein kinase
Ang II: angiotensin II
AT1: angiotensin 1 receptor
AT2: angiotensin 2 receptor
ATP: adenosine triphosphate
BCA: bicinchoninic acid
BSA: bovine serum albumin
CABG: coronary artery bypass graft
CAD: coronary artery disease
CVD: cardiovascular disease
DMEM: Dulbecco’s modified Eagle medium
DMSO: dimethyl sulfoxide
DNA: deoxyribonucleic acid
DPBS: Dulbecco’s phosphate buffered saline
ECAR: extracellular acidification rate
EDTA: ethylenediaminetetraacetic acid
ERK: extracellular signal-regulated kinase
FCCP: Carbonyl cyanide p-(trifluoromethoxy)phenylhydrazone
GAPDH: glyceraldehyde-3-phosphate dehydrogenase
HSVSMC: human saphenous vein smooth muscle cell
HUVEC: human umbilical vein endothelial cell
IL-6: interleukin-6
JAK: Janus kinase
MAPK: mitogen-activated protein kinase
NAD: nicotinamide adenine dinucleotide
NIH: neointimal hyperplasia
NOS: nitric oxide synthase
OCR: oxygen consumption rate
PAH: pulmonary arterial hypertension
PBS: phosphate-buffered saline
PBST: phosphate-buffered saline with Tween® detergent
PDGF: platelet-derived growth factor
PH: pulmonary hypertension
sIL-6Rα: soluble interleukin-6 receptor
STAT: signal transducer and activator of transcription
T2DM: type 2 diabetes mellitus
TGF: transforming growth factor
VEC: vascular endothelial cell
VGF: vein graft failure
VSMC: vascular smooth muscle cell.

## Authors’ contribution

**IOB:** Conceptualization, Methodology, Investigation, Formal analysis, Visualization, Funding acquisition, Writing – original draft. **FTM**: Methodology, Investigation, Formal analysis. **GAD:** Investigation, Visualization, Writing – review and editing. **JPH:** Writing – ethics and project administration. **ML:** Investigation, Validation, and Writing – review & editing. **KR-S: Supervision,** Validation, Writing – review & editing. **RGS: Supervision,** Validation, Writing – review & editing. **TMP:** Conceptualization, Supervision, Methodology, Project administration, Funding acquisition, Writing – review & editing.

## Data availability

Data will be provided upon request.

