## Supplementary Material for "JAK and MEK Pathways as Therapeutic Targets for Saphenous Vein Smooth Muscle Cell Dysfunction in Type 2 Diabetes *via* Regulation of Mitochondrial Activity"

**SUPPLEMENTARY DATA**

1. **Materials**

| **Company/Supplier** | **Catalogue number** |
| --- | --- |
| **Abcam, Cambridge, UK** |  |
| Ruxolitinib | ab141356 |
| Oligomycin | ab141829 |
| Carbonyl cyanide p-(trifluoromethoxy) phenylhydrazone (FCCP) | ab120081 |
| Rotenone | ab143145 |
| Antimycin A | ab141904 |
| Anti-STAT3 antibody (EPR787Y) | ab68153 |
| **Agilent Technologies Incorporated, Santa Clara, USA** |  |
| Seahorse XF FluxPaks | 103022-100 |
| XF DMEM Medium, pH 7.4, 500ml, with 5 mm HEPES, without phenol red, sodium bicarbonate, glucose, sodium pyruvate, L-glutamine | 103575-100 |
| **Cell Signalling Technology, Massachusetts, USA** |  |
| Phospho-Stat3 (Tyr705) antibody | 9131 |
| Phospho-p44/42 MAPK (Erk1/2) (Thr202/Tyr204) (E10) mouse antibody | 9106L |
| p44/42 MAPK (Erk1/2) antibody | 9102L |
| **PromoCell GmbH, Heidelberg, Germany** |  |
| Smooth muscle cell growth medium 2 kit | C-22162 |
| **R&D systems, Minnesota, USA** |  |
| Recombinant human IL-6 protein | 206-IL |
| Recombinant human IL-6 R alpha protein | 227-SR-025 |
| **Sigma-Aldrich (Merck), Dorset, UK** |  |
| Ammonium persulfate (APS) | A3678 |
| Anti-O-GlcNAc Transferase (DM-17) antibody produced in rabbit | O6264 |
| Hepes | H3375 |
| Fetal bovine serum | F7524 |
| Angiotensin II human | A9525-5mg |
| Thrombin from bovine plasma | T4648-1ku |
| **Stratech, Cambridgeshire, UK** |  |
| Trametinib | GSK1120212 |
| **Thermo Scientific, Massachusetts, USA** |  |
| RECOMBINANT HUMAN PDGF-BB | 10531285 |

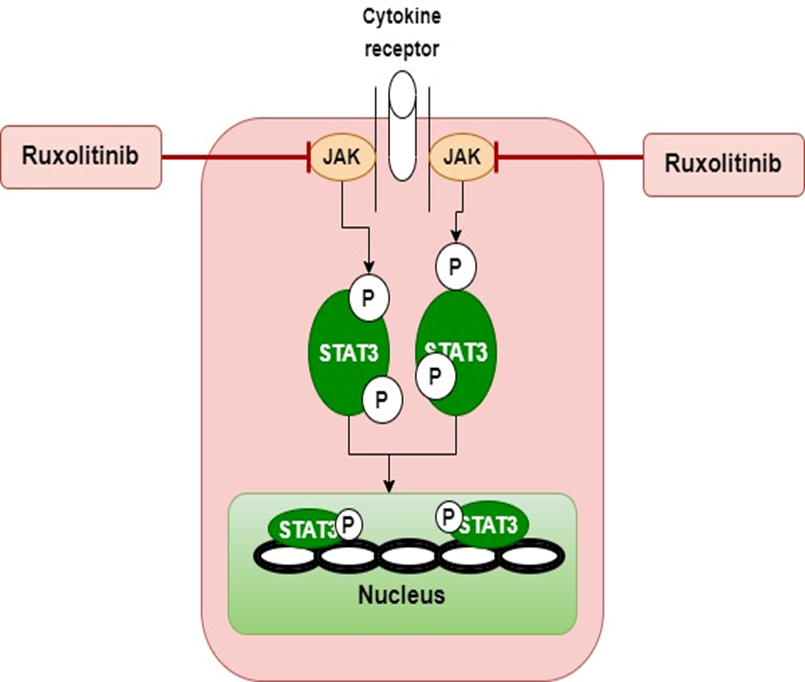

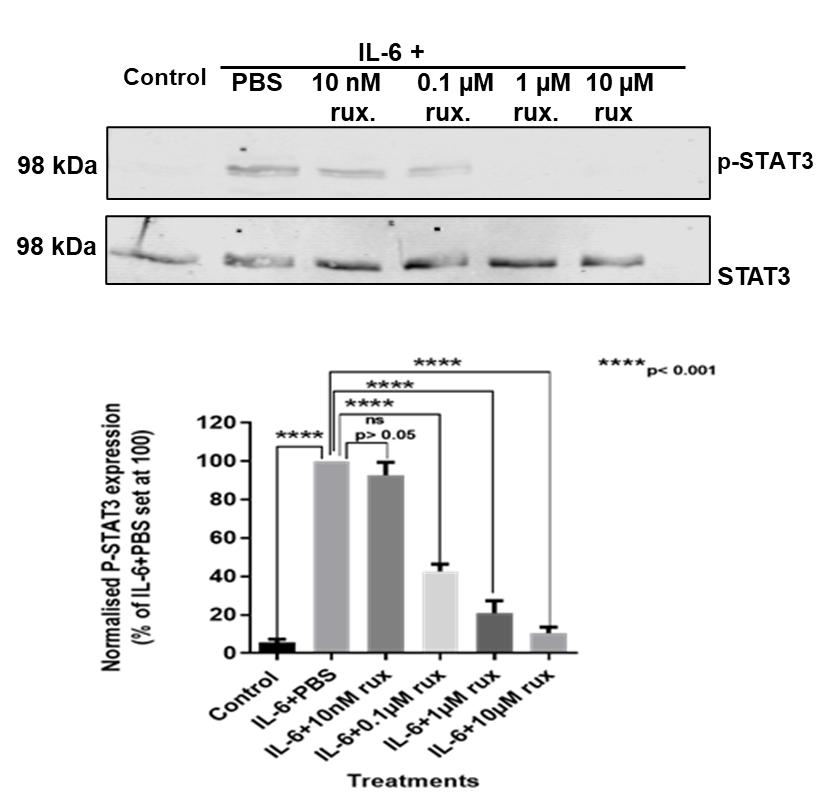

**B**

**C**

**A**

**Figure S1: Downstream activation and inhibition of the JAK/STAT signalling pathway. (A)** Schematic of downstream activation and inhibition of the JAK/STAT signalling pathway. The JAK/STAT pathway transmits information from chemical signals outside of a cell to the cell nucleus where it activates genes through transcription. This is mediated by activation of receptors such as IL-6R and proteins, Janus kinases (JAKs), and signal transducer and activator of transcription (STAT). Ruxolitinib (JAK 1/2-selective inhibitor) blocks this signalling pathway.

**(B)** Representative western blot of downstream inhibition of IL-6/sIL-6Rα-stimulated JAK1/2-mediated phosphorylation of STAT3 (98kDa) on Tyr705 by ruxolitinib. Lower panel: expression of total STAT3 (**C)** Densitometric analysis of p-STAT3 normalised to total STAT3. Data are expressed as mean+/-SEM and from n=4 biological replicates using HSVSMC samples from different non-diabetic patients. Statistical significance was assessed using one-way ANOVA followed by Dunnett’s post-hoc test to determine significant differences between means.IL-6: IL-6/sIL-6Rα; PBS: Phosphate-buffered saline; rux: Ruxolitinib.

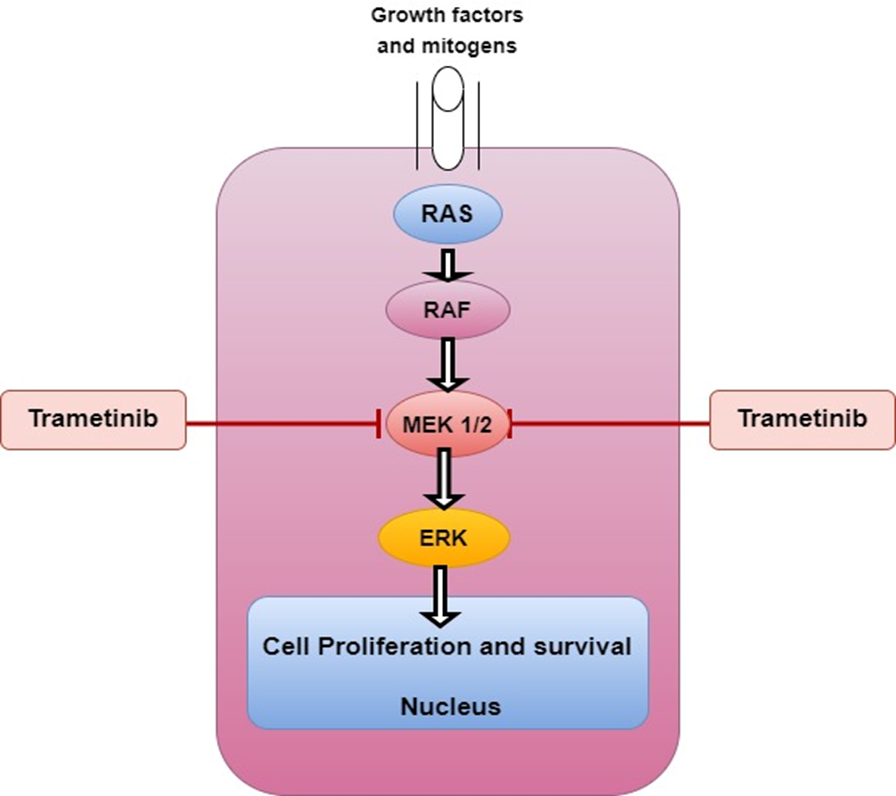

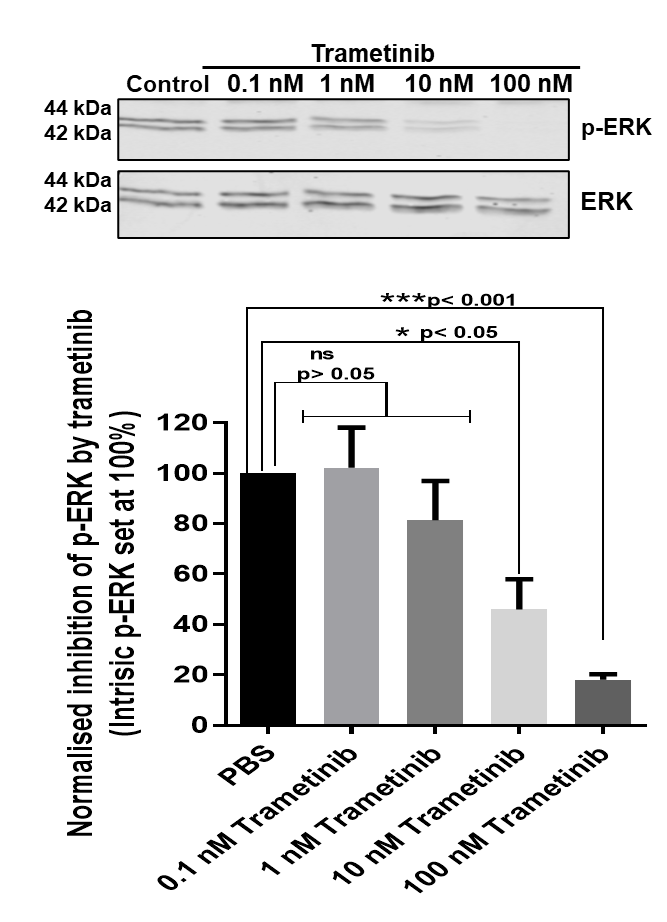

**C**

**B**

**A**

**Figure S2: Downstream activation and inhibition of the MAPK/ERK signalling pathway.** **(A)** Schematic of downstream activation and inhibition of the MEK/ERK signalling pathway. Mitogens trigger activation of the small GTPase Ras protein, which in turn activates the Ser/Thr-directed protein kinase Raf, which phosphorylates and activates MEK 1/2. MEK1/2 and then phosphorylate and stimulate ERK1,2 (Thr202/185 and Tyr204/187 in human ERK1/2 respectively). ERK further activates a panel of substrates that promote cell proliferation. **(B)** Upper panel: representative western blot of downstream inhibition of the MEK 1/2 by trametinib. Lower panel: expression of total ERK (44/42 kDa). **(C)** Densitometric analysis of p-ERK at both 44/42 kDa normalised to total ERK at both 44/42 kDa. Data are expressed as mean+/-SEM and from n=4 biological replicates using HSVSMC samples from different non-diabetic patients. Statistical significance was assessed using one-way ANOVA followed by Dunnett’s post-hoc test to determine significant differences between means. PBS: Phosphate-buffered saline.

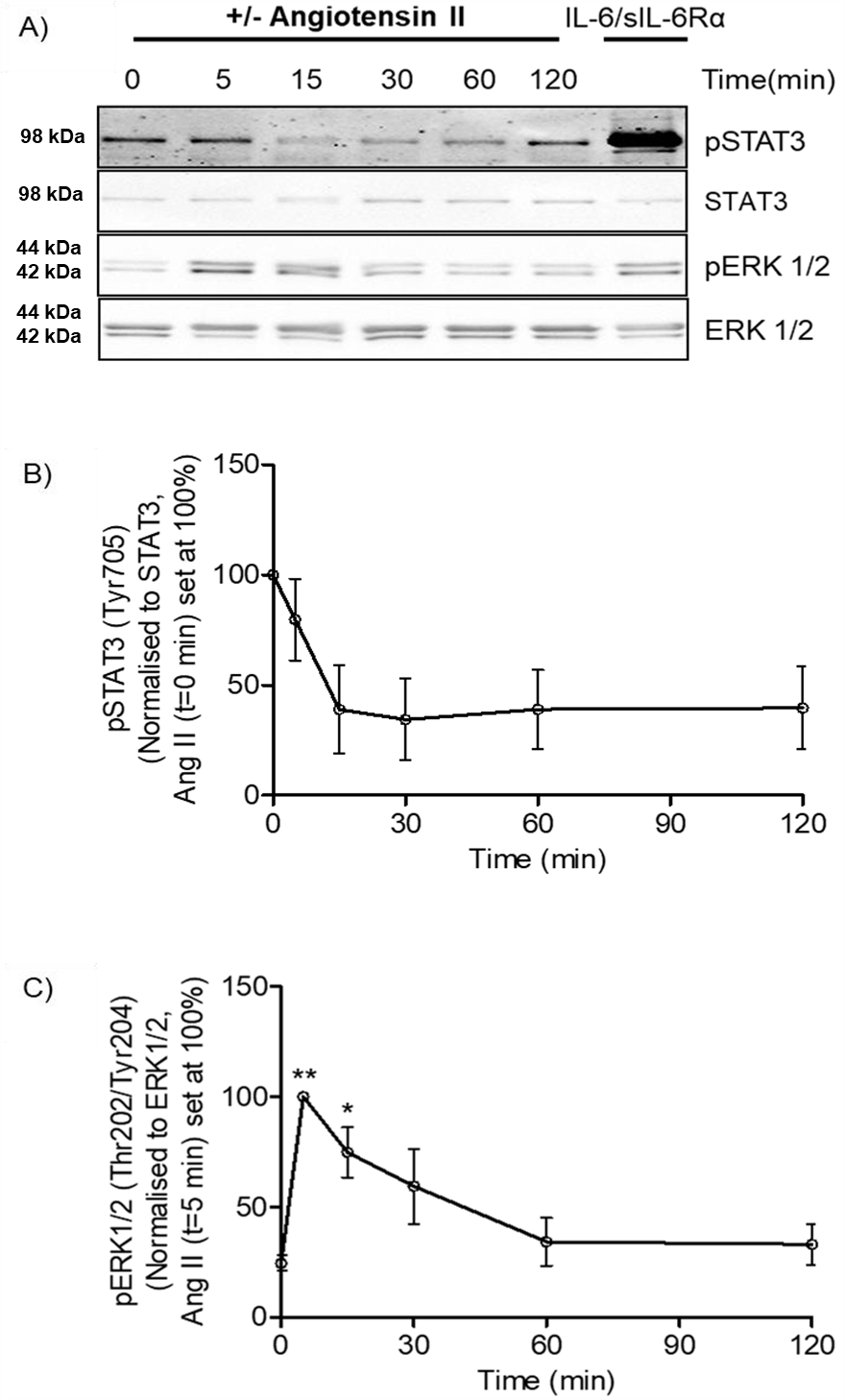

**Figure S3:** Time course of the effects of angiotensin II treatment on STAT3 and ERK1/2 phosphorylation in HSVSMCs. HSVSMCs were either treated with vehicle or Angiotensin II (Ang II) (0.1µM) for the indicated times or sIL-6Rα/IL-6 for 30 minutes before cell harvest. **(A)** Protein-equalised cell lysates were fractionated by SDS-PAGE for immunoblotting. P-STAT3 (Tyr705) **(B)** and P ERK1/2 (Thr202/Tyr204) **(C)** levels were normalised to total STAT3 and ERK1/2, respectively, and measured relative to the maximum response (set at 100%). Data are presented as mean ± S.E.M. Data are expressed as mean+/-SEM and from n=4 biological replicates using HSVSMC samples from different non-diabetic patients. Statistical significance was assessed using one-way ANOVA followed by Dunnett’s post-hoc test to determine significant differences between means, ******P* < 0.05; ***P* < 0.01 versus t=0 minutes (min).
